# Investigating genetic diversity within the most abundant and prevalent non-pathogenic leaf-associated bacteria interacting with *Arabidopsis thaliana* in natural habitats

**DOI:** 10.1101/2022.07.05.498547

**Authors:** Daniela Ramirez-Sanchez, Chrystel Gibelin-Viala, Baptiste Mayjonade, Rémi Duflos, Elodie Belmonte, Vincent Pailler, Claudia Bartoli, Sébastien Carrere, Fabienne Vailleau, Fabrice Roux, Fabienne Vailleau

## Abstract

Plants interact simultaneously with highly diversified microbes defined as the plant microbiota. Microbiota modulates plant health and appears as a promising lever to develop innovative, sustainable and eco-friendly agro-ecosystems. Key patterns of microbiota assemblages in plants have been revealed by an extensive number of studies based on taxonomic profiling by metabarcoding. However, understanding the functionality of microbiota and identifying the genetic and molecular mechanisms underlying the interplay between plants and their microbiota are still in its infancy and relies on reductionist approaches primarily based on the establishment of representative microbial collections. In *Arabidopsis thaliana*, most of these microbial collections include one strain per OTU isolated from a limited number of habitats, thereby neglecting the ecological potential of genetic diversity within microbial species to affect the plant-microbiota molecular dialog. With this study, we aimed at estimating the extent of genetic variation between strains within the most abundant and prevalent leaf-associated non-pathogenic bacterial species in 163 natural populations of *A. thaliana* located south-west of France. By combining a culture-based collection approach consisting of the isolation of more than 7,000 bacterial colonies with an informative-driven approach, we isolated 35 pure strains from eight non-pathogenic bacterial species. We detected significant intra-specific genetic variation at the genomic level and for growth rate in synthetic media. In addition, significant host genetic variation was detected in response to most bacterial strains in *in vitro* conditions, with the presence of both negative and positive responses on plant growth. Our study provides new genetic and genomic resources for a better understanding of the plant-microbe ecological interactions at the microbiota level. We also highlight the need of considering genetic variation in both non-pathogenic bacterial species and *A. thaliana* to decipher the genetic and molecular mechanisms involved in the ecologically relevant dialog between hosts and leaf microbiota.

## INTRODUCTION

During their life cycle, plants interact simultaneously with a large and diverse set of microbial organisms – viruses, bacteria, fungi, oomycetes, archaea and protists – defined as the plant microbiota and often referred as the second genome of plants, in agreement with the holobiont/hologenome theory of evolution (Roux and Bergelson, 2016; Morris, 2018). Plant microbiota mainly originates from the soil reservoirs, even if a non-negligible fraction of microbes may originate from the aerial sphere (Müller et al., 2016). For long, microbes were considered harmful to the plants. Accordingly, an extensive knowledge has been acquired on the molecular dialog between plants and pathogens (Jones and Dangl, 2006; Deslandes and Rivas, 2012; Roux et al., 2014; Uhse and Djamei, 2018; Delplace et al., 2021; Ngou et al., 2022). Nowadays, it is well-known that a large fraction of plant-associated microbes are beneficial for their hosts. For instance, plant growth promoting bacteria (PGPB) can mobilize and provision nutrients to plants either directly or indirectly by enhancing interaction with other microbes. For example, in wheat, rhizobia strains can both increase wheat root development and improve the wheat-arbuscular mycorrhiza interactions (Bartoli et al., 2020). Beneficial bacteria can also provide protection against pathogens, either directly through the production of antimicrobial compounds, competition for nutrients and colonization sites, or indirectly with the induction of plant defense such as systemic resistance mechanisms (Berendsen et al., 2012; Bulgarelli et al., 2013; Pieterse et al., 2014; Trivedi et al., 2020; Glick and Gamalero, 2021).

During the last decade, the rise of next-generation sequencing technologies enabled a deep taxonomic characterization of microbial communities of the rhizosphere, root and leaf compartments in diverse wild and crop species, which in turn allowed to identify key patterns of plant microbiota assemblages (Müller et al., 2016). Firstly, microbial communities are not randomly assembled (Bulgarelli et al., 2012; Lundberg et al., 2012). In line with this, strong and reproducible successional dynamics were observed in phyllosphere communities (Maignien et al., 2014), which are mainly colonized by bacterial members of the three phyla Proteobacteria, Actinobacteria and Bacteroidetes (Vorholt, 2012; Bodenhausen et al., 2013). Secondly, microbiota composition and assemblage largely differ between plant compartments (Coleman-Derr et al., 2016; Wagner et al., 2016; Trivedi et al., 2020), with phyllosphere communities being relatively less diverse and presenting more inter-individual variability than rhizosphere communities (Lebeis, 2015). Microbial communities also largely differ between endophytic and epiphytic compartments, with microbiota richness in roots being higher in the episphere than in the endosphere, while the reverse is true for leaves (Bodenhausen et al., 2013; Lebeis et al., 2015). Thirdly, extensive variation in microbiota composition was detected *in situ* among natural populations at diverse geographical scales (Bodenhausen et al., 2013; Agler et al., 2016; Coleman-Derr et al., 2016; Geremia et al., 2016; Thiergart et al., 2020). For instance, at a regional scale, the level of differentiation for bacterial communities among more than 160 natural populations of *Arabidopsis thaliana* reached up to ∼81% in both leaf and root compartments (Bartoli et al., 2018). Spatial variation in microbiota composition results from the intermixed effects of host genetics and environmental variation, such as climate, soil agronomic properties and companion plant species (Aleklett et al., 2015; Geremia et al., 2016; Thiergart et al., 2020; Meyer et al., 2022). Fourthly, both artificial and natural genetic variation indicate that variation of host control within plant species is governed by many genes of small effect, with the exception of genes involved in the circadian clock or the biosynthesis of specialized metabolites such as thalianin and arabidin (Bergelson et al., 2021).

Taxonomic profiling of microbial communities were then followed, albeit only in few instances, by studies dedicated to functionally characterize plant-microbiota interactions. Plant-microbiota functional characterization has been mostly conducted in *A. thaliana* and first required the establishment of representative microbial collections. Starting with a collection of microbes isolated from *A. thaliana* plants grown on an agricultural soil from Germany in greenhouse conditions (Bai et al., 2015; Durán et al., 2018), bacterial root commensals were demonstrated to both deeply influence fungal and oomycetal community structure and protect plant against root-derived filamentous eukaryotes (Durán et al., 2018). Using synthetic communities (SynCom) derived from the same collection and inoculated on germ-free plants, recolonization experiments demonstrated specialization of bacteria to their respective plant niche from which they originate (*i.e.* leaf and root), while forming assemblies resembling natural bacterial communities (Bai et al., 2015). Still based on the same collection, a SynCom approach also revealed that community assemblies in phyllosphere are historically contingent and subject to priority effects (Carlström et al., 2019). On the other hand, a collection of bacterial isolates from an agricultural soil in USA highlighted the close interplay of root microbiota with plant immunity (Lebeis et al., 2015; Teixeira et al., 2021) and plant response to nutritional stress (Castrillo et al., 2017). Finally, when bacterial collections are paired with genome sequencing, a large overlap of genome-encoded functional capabilities was revealed between leaf- and root-derived bacteria (Bai et al., 2015).

While these studies started to provide a glimpse of the functional mechanisms underlying host-microbiota dialog, most microbial collections from *A. thaliana* were established on the isolation of one representative strain per Operational Taxonomic Unit (OTU) and from a very limited number of agricultural and natural sites, thereby neglecting the ecological potential of genetic diversity within microbial species to strongly impact the outcomes of plant-microbiota interactions. Yet, extensive genetic variation among natural populations of *A. thaliana* was detected for virulence among strains of the main bacterial species of its pathobiota (*i.e. Pseudomonas syringae*, *Pseudomonas viridiflava* and *Xanthomonas campestris*) (Bartoli et al., 2018), which is in line with the large genomic diversity observed in both *P. syringae* and *Xanthomonas arboricola* at a local population scale (Karasov et al., 2014, 2018; Wang et al., 2018).

As a first step to investigate the effects of genetic diversity within microbial species on plant-microbiota interactions and to identify the underlying genetic and molecular mechanisms in both host plant and microbes, we aimed at establishing an informative collection of several isolates of the 12 most abundant and prevalent non-pathogenic leaf-associated bacterial OTUs among 163 natural populations of *A. thaliana* located south-west of France (Bartoli et al., 2018). We focused on bacterial communities in the phyllosphere because of (i) the 60% contribution of the phyllosphere to the biomass across all taxa on Earth, (ii) to the complex natural habitats provided by leaves with extensive microscale variation and diverse nutriments provided for bacterial growth, and (iii) the benefits provided by phyllosphere bacteria to their hosts through production of plant growth-promoting hormones and protection against pathogen infection (Lindow and Brandl, 2003; Vorholt, 2012; Maignien et al., 2014; Koskella, 2020). We first describe the isolation of several strains of seven of the most abundant and prevalent bacterial OTUs by combining a community-based culture (CBC) approach based on the amplification of a fraction of the *gyrB* gene (Bartoli et al., 2018), with an informative-driven approach based on specific culture media and the design of specific primers for each OTU. Based on 22 representative bacterial strains, we then report the extent of (i) genomic variation among and within OTUs, (ii) genetic variation on *in vitro* bacterial growth kinetics, and (iii) genotype-by-genotype (GxG) interactions between the isolates and eight genotypes of *A. thaliana* (including seven local accessions and the reference genotype Col-0) on plant growth when inoculated at the seed and seedling stages.

## MATERIAL AND METHODS

### Identification of the most prevalent and abundant OTUs

One hundred and sixty-three natural populations of *A. thaliana* located south-west of France have been previously characterized for bacterial communities by a metabarcoding approach based on a portion of the *gyrB* gene (Watanabe et al., 2001; Bartoli et al., 2018). More precisely, 821 rosettes collected *in situ* in autumn 2014 and spring 2015 were characterized. For each rosette sample, we estimated the relative abundance of each of the 6,627 most abundant OTUs out of the 278,333 identified OTUs (Bartoli et al., 2018). Then, we selected OTUs both present in more than 5% of rosettes (n > 41) and with a mean relative abundance across all rosettes above 0.7%, resulting in 13 leaf OTUs. From these 13 OTUs, we removed OTU8 (prevalence = 6.7%, mean relative abundance = 1.1%) corresponding to *P. viridiflava* for which 74 isolated strains from the natural populations of *A. thaliana* were confirmed to be pathogenic (Bartoli et al., 2018). We therefore focused on the remaining 12 OTUs (Figure 1A) for which taxonomic affiliation based on *gyrB* amplicon was identified according to GenBank BLASTN (Supplementary Data Set 1).

**Figure 1.**
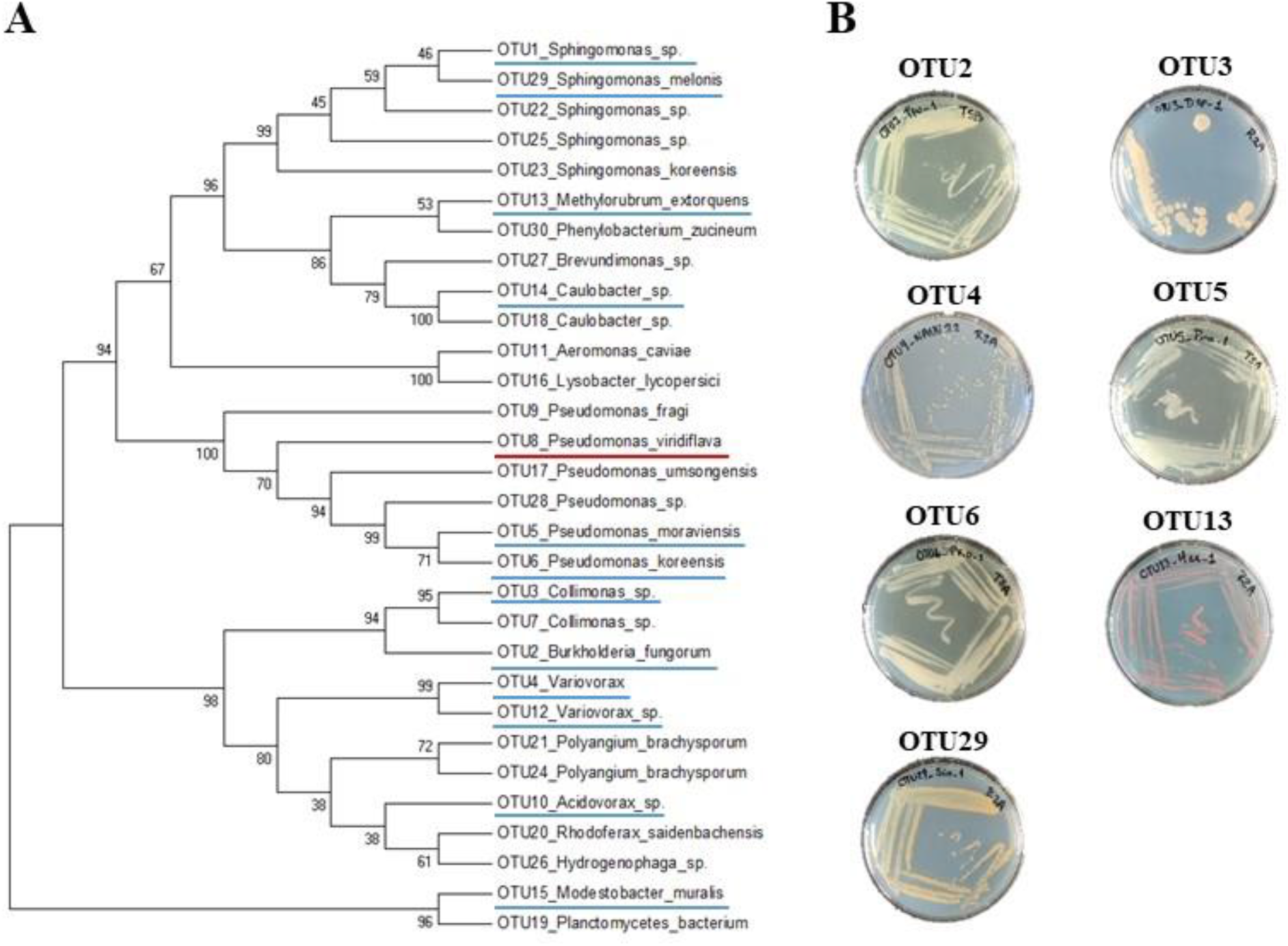
Phylogenetic and morphological diversity of the 12 most prevalent and abundant OTUs of the leaf compartment among 163 natural populations of *A. thaliana* located south-west of France. **(A)** *gyrB* amplicon based phylogenetic tree of the 30 most abundant OTUs of the leaf compartment among 163 natural populations of *A. thaliana*. The 12 most prevalent and abundant OTUs are underlined in blue. The OTU8 corresponding to *Pseudomonas viridiflava* for which 74 isolated strains were confirmed to be pathogenic, is underlined in red. **(B)** Morphological diversity of the seven OTUs for which representative strains have been isolated.

### Isolation of strains

#### Leaf samples

To isolate bacterial strains associated with the selected 12 OTUs, a total of 205 and 300 plants were collected across 91 populations in autumn 2014 and 152 populations in spring 2015, respectively. Plants were excavated using flame-sterilized spoons and then manipulated with flame-sterilized forceps on a sterilized porcelain plate. Gloves and plate were sterilized by using Surface’SafeAnios®. Rosettes were rinsed into individual tubes filled with sterilized distilled water to remove all visible dirt. Rosettes were then placed into sterilized tubes and immediately stored in a cooler filled with ice. The 505 rosettes were ground in 500 μl of sterile distilled water solution and stored in a 20% glycerol solution at -80°C. The epiphytic and endophytic compartments of rosettes were not separated.

#### Community-based culture approach

In order to isolate multiple strains for each of the 12 candidate OTUs, 48 strains were randomly isolated from two rosettes (*i.e.* 24 strains per rosette) for each of the 152 natural populations sampled in spring 2015 (Supplementary Figure 1, Supplementary Table 1). To do so, serial dilutions of glycerol stocks were performed from 10^-2^ to 10^-7^ using sterile distilled water. For each of three successive serial dilutions (10^-5^ to 10^-7^), 100 µl were plated on a tryptic soy broth (TSB) agar medium (3 g/L TSB and 12 g/L agar) in 10 cm ×10cm square Petri dishes. After 48H of growth at 28°C, 24 colonies were randomly chosen for each rosette. Each colony was picked with a sterile 10 µl tip attached to a P10 pipette, then dipped in a well of a 96-well PCR plate filled with 20 µl of a *gyrB* PCR mix (4 µl of 5x GoTaq buffer (Promega, Madison, WI, USA), 0.4 µl of 10mM dNTP, 0.4 µl of each 10µM primer, 0.1 µl of Tween 20, 0.2 µl of 5u/µl GoTaq G2 Polymerase (Promega), and 14.5 µl of PCR-grade water) and mixed by pipetting up and down 3 times. Afterwards, the same tip was dipped and mixed by pipetting up and down 3 times in a well of a 96-well 0.8 ml deep well plate filled with 300 μl liquid 3g/L TSB medium. After incubation at 28°C for 48H, 150 µl of 60% glycerol was added and plates were stored at -80°C. Amplification was then performed as follows: 94°C (2 min) followed by 40 cycles of amplification at 94°C (30 s), 50°C (60 s), and 68°C (90 s), with a final extension step of 10 min at 68°C. PCR products were checked on a 2% agarose gel.

In order to multiplex the 7,259 amplicons, corresponding to 7,259 picked colonies, on a single Miseq run, seven forward and seven reverse *gyrB* primers containing different internal tags were combined for each PCR plate (Supplementary Data Set 2). The 76 96-well PCR plates were then pooled into four pools of 96 samples. Pooled amplicons were purified by using Clean PCR beads (CleanNA, Waddinxveen, The Netherlands) and following the manufacturer’s instructions. Purified amplicons were quantified with a Nanodrop and appropriately diluted to obtain an equimolar concentration. The second PCR was prepared in 50 µl and contained 2 µl of equimolar PCR-purified products, 5 µl of 10X MTP Taq buffer (Sigma-Aldrich, Saint-Louis, MO, USA), 1 µl of 10 mM dNTP, 1.25 µl of 20 µM forward P5 primer, 1.25 µl of 20µM barcoded reverse P7 primer, 0.5 µl MTP Taq DNA polymerase (Sigma-Aldrich), and 39 µl of PCR-grade water. The list and sequences of the primers are given in the Supplementary Data Set 2. Amplification was performed as follow: 94°C (60 s) followed by 12 cycles of 94°C (60 s), 65°C (60 s), 72°C (60 s) and finally 72°C (10 min). Amplicons were further purified and quantified as described above to obtain a unique equimolar pool. The latter was quantified by real-time quantitative reverse transcriptase-polymerase chain reaction and then sequenced on a Illumina MiSeq system with 2 × 250bp paired-end reads (Illumina Inc., San Diego, CA, USA) at the GeT-PlaGe Platform (Toulouse, France). MS-102-3003 MiSeq Reagent Kit v3 600 cycle was used for this purpose.

Demultiplexed Illumina barcoded libraries (available at NCBI SRA under SRP372434 accession) were trimmed with “fastx_trimmer-t 51” (http://hannonlab.cshl.edu/fastx_toolkit/, access on January 6^th^ 2022) to produce 200nt high-quality reads and assembled using FLASH (Magoč and Salzberg, 2011) assembler (version 1.2.11) with the following parameters “--max-overlap 120 --min-overlap 50”. Assembled amplicons were then scanned to identify internal tags on both ends using exact match regular expressions (Supplementary Data Set 3, Supplementary Data Set 4).

A total of 14,956,927 reads were obtained for 7,066 out of the 7,259 sequenced colonies, with a mean number of reads per sample of 2,117 (confidence intervals 95%: 177 – 6889). To taxonomically classify these 7,066 samples, we used the DADA2 (Callahan et al., 2016) pipeline (version 1.6.0) to identify 6,362 amplicon sequence variants (ASVs) that we assigned to the closest taxon using a manually curated *gyrB* database composed of 38,929 sequences (version 2) (Supplementary Data Set 5) (Barret et al., 2015).

Among the 7,066 samples, we identified 5 samples with at least 10% of the reads belonging to an ASV matching to OTU2 with an identity ≥ 98%. With the same criteria, we identified 55 samples for OTU3, 21 samples for OTU5, 365 samples for OTU6 and 59 samples for OTU29.

#### Informative-driven approach to isolate pure strains

To isolate pure strains from the 505 identified samples, we adopted an approach combining the use of selective medium including specific antibiotics and genus-specific primers (Supplementary Figure 1, Supplementary Table 1, Supplementary Table 2). For OTUs missing specific primers in the literature, we designed specific ones based on *gyrB* sequences (Supplementary Material, Supplementary Table 3). With this combined approach, we retrieved 3 pure strains for OTU2, 7 pure strains for OTU3, 8 pure strains for OTU5, 6 pure strains for OTU6 and 8 pure strains for OTU29.

For the isolation of pure strains for the remaining seven OTUs (OTU1, OTU4, OTU10, OTU12, OTU13, OTU14 and OTU15), the same strategy was applied on rosette samples collected in either autumn or spring, from natural populations presenting the highest relative abundance for these OTUs in their leaf microbiota (Bartoli et al., 2018). Serial dilutions from glycerol stocks were performed from 10^-2^ to 10^-7^ using sterile distillated water. For each dilution, 100 µl were plated on the corresponding selective medium in 10 cm ×10 cm square Petri dishes. Colonies were grown at 28°C for at least 48H. Colonies selected based on morphological aspects and for amplifying with the specific primers, were further amplified and sequenced with the Sanger technology for the *gyrB* gene with universal primers (Barret et al., 2015). The resulting *gyr*B sequences were blasted against both (i) a database that contains the sequences of the 6,627 most abundant OTUs identified across our 163 natural populations of *A. thaliana* (Bartoli et al., 2018), and (ii) the NCBI database. All the bacterial strains with a *gyrB* sequence matching one of the *gyrB* OTU sequences of interest with an identity ≥ 98% and over ≥ 200 bp were stored in a 20% glycerol solution at -80°C. After using this approach on the remaining seven OTUs, we retrieved one pure strain for OTU4 and two pure strains for OTU13.

Finally, by combining the CBC and informative-driven approaches, we isolated a total of 35 pure strains for seven OTUs (OTU2, OUT3, OTU4, OTU5, OTU6, OTU13 and OTU29).

### Genomic sequences

For a subset of 22 strains maximizing the geographical diversity among the 35 pure isolated strains (Table 1, Supplementary Table 1), high molecular weight DNA was extracted from bacterial culture grown for 24H at 28°C in appropriate media (TSB for OTU2, R2A for OTU3, OTU4, OTU13 and TSA for OTU5, OTU6, OTU29) following the protocols described in (Mayjonade et al., 2016). The purity of the DNA samples was assessed on a NanoDrop spectrophotometer (A260/A280 and A260/A230 ratios) and the DNA concentration was measured with a Qubit dsDNA HS reagent Assay Kit (Life Technologies). Single-molecule Real-time long reads sequencing was performed at the Gentyane core facility (Clermont-Ferrand, France) with a PacBio Sequel II system (Pacific Biosciences, Menlo Park, CA, USA). The SMRTBell library was prepared using a SMRTbell Express 2 Template prep kit, following the “procedure and checklist -preparing Multiplexed Microbial Libraries using SMRTbell Express Template prep kit 2.0” protocol. Briefly, for each strain, 1 µg of genomic DNA was sheared into approximatively 10 kb fragments using g-tubes (Covaris, England). A Fragment Analyzer (Agilent Technologies, Santa Clara, CA, USA) assay was used to assess the size distribution of fragments. Sheared genomic DNA samples were carried into the enzymatic reactions to remove the single-strand overhangs and to repair any damage that may be present on the DNA backbone. An A-tailing reaction followed by the overhang adapter ligation (SMRTbell Barcoded Adapter Plate 3.0) was conducted to generate the SMRT Bell templates. The samples were then purified with 0.45X AMPure PB Beads to obtain the final libraries at around 10 kb. The SMRTBell libraries were checked for quality using a Fragment Analyzer (Agilent Technologies) and quantified with a Qubit dsDNA HS Assay Kit. A ready-to-sequence SMRTBell Polymerase Complex was created using a Binding Kit 2.2 (PacBio) and the primer V5. The adaptive loading protocol was used according to the manufacturer’s instructions. The PacBio Sequel instrument was programmed to load a 90 pM library and sequenced in CCS mode on a PacBio SMRTcell 8M, with the Sequencing Plate 2.0 (Pacific Biosciences), 2 hours of pre-extension time and acquiring one movie of 15 hours.

**Table 1.** List of the 12 most prevalent and abundant OTUs of the leaf compartment among 163 natural populations of *A. thaliana* located south-west of France. Lines highlighted in light grey correspond to OTUs for which representative strains have been isolated.

| OTU<br>Bartoli *et al.* (2018) | *gyrB* based taxonomic affiliation | Prevalence (%) | Mean relative abundance (%) | ANI based taxonomic affiliation | Strain name |
| --- | --- | --- | --- | --- | --- |
| OTU_1 | *Sphingomonas* sp. | 44.6 | 16.07 |  |  |
| OTU_2 | *Paraburkholderia fungorum* | 5.7 | 1.79 | *Paraburkholderia fungorum* | OTU2_Pfu_1<br>OTU2_Pfu_2<br>OTU2_Pfu_3 |
| OTU_3 | *Collimonas* sp. | 43.1 | 5.65 | *Oxalobacteraceae* bacterium | OTU3a_Oxa_1<br>OTU3a_Oxa_2<br>OTU3b_Oxa_1<br>OTU3b_Oxa_2 |
| OTU_4 | *Variovorax* sp. | 41.8 | 5.33 | *Comamonadaceae* bacterium | OTU4_Com_1 |
| OTU_5 | *Pseudomonas moraviensis* | 12.3 | 1.18 | *Pseudomonas moraviensis* | OTU5_Pmo_1<br>OTU5_Pmo_2<br>OTU5_Pmo_3 |
| OTU_6 | *Pseudomonas koreensis* | 12.2 | 1.11 | *Pseudomonas siliginis* | OTU6_Psi_1<br>OTU6_Psi_2<br>OTU6_Psi_3<br>OTU6_Psi_4<br>OTU6_Psi_5<br>OTU6_Psi_6 |
| OTU_10 | *Acidovorax* sp. | 11.9 | 1.36 |  | - |
| OTU_12 | *Variovorax* sp. | 9.4 | 0.7 |  | - |
| OTU_13 | *Methylobacterium extorquens* | 14.1 | 0.98 | *Methylobacterium* sp. | OTU13_Msp_1<br>OTU13_Msp_2 |
| OTU_14 | *Caulobacter* sp. | 9.9 | 0.89 |  | - |
| OTU_15 | *Modestobacter* sp. | 9.7 | 0.85 |  | - |
| OTU_29 | *Sphingomonas melonis* | 14 | 0.89 | *Sphingomonadaceae* bacterium | OTU29_Sph_1<br>OTU29_Sph_2<br>OTU29_Sph_3 |

For each strain, raw PacBio sequences were self-corrected using ccs software (version 6.0.0) with a minimum of 3 passes. Assembly was performed with FLYE assembler (version 2.9) (Kolmogorov et al., 2020) with the specific parameters “--pacbio-hifi” and for each strain an estimated genome assembly size (from 4Mb to 9Mb). We used *DnaA* gene sequence to determine chromosome start for each assembled circular chromosome and annotate genomes using bacannot (version 3.1) (de Almeida and Pappas Jr, 2022), InterProScan (version 5.54-87.0) (Jones et al., 2014) and Blast2GO (version 1.5.1) (Götz et al., 2008) to infer putative functions.

### ANI taxonomic affiliation

In order to accurately assign the taxonomy of the sequenced strains, we used FastANI, an algorithm representing the Average Nucleotide Identity (ANI) of all orthologous genes shared between two genomes (Jain et al., 2018). To do so, for each OTU, the 10 closest genomes were retrieved from each of three quality-controlled and curated databases, *i.e.* JSpecies (Richter et al., 2016), GTDB-Tk (Chaumeil et al., 2019) and TYGS (Meier-Kolthoff and Göker, 2019). Identical genomes among the three databases were considered only once. Subsequently, FastANI was used to compute pairwise ANI between each strain and the JSpecies-GTDB-Tk-TYGS list of closest genomes. A strain was assigned to a species if the ANI value was > 95%. To confirm that all strains from a given OTU belong to the same bacterial species, pairwise ANI values among strains were estimated.

### Orthogroups

We looked for orthologous and paralogous genes using OrthoFinder (Emms and Kelly, 2015) software (version 2.5.4, default parameters with diamond software as aligner) in order to extract core-genes. OrthoFinder results are available and queryable through Family-Companion (Cottret et al., 2018) software at this url: https://doi.org/10.25794/e29x-aq46. Based on Orthofinder results, a list of orthogroups was extracted for each of the six strains belonging to the OTU6. To estimate and illustrate the number of common and specific orthogroups between these six strains, a Venn diagram was generated using jvenn (Bardou et al., 2014).

### Growth kinetics

The 22 selected strains were streaked onto their respective medium (Supplementary Table 1). All strains were grown at 28°C during 2 days (OTU2, OTU3, OTU4, OTU5 and OTU6) or 7 days (OTU13 and OTU29). Colonies of each strain were then scrapped off with an inoculating loop and re-suspended in 500 µl of sterile distilled water. 250 µl of that volume were sprayed in a new plate and incubated at 28°C overnight. Fresh colonies were scrapped off with an inoculating loop and diluted in 1 ml of sterile water. The suspension was adjusted to an OD600nm of 0.15 in a volume of 200 µl deposited in a 96-well Greiner plate. For each strain, monitoring was performed both in the R2A minimal nutrient medium and the TSB rich nutrient medium. Bacterial growth was monitored using a microplate spectrophotometer (FLUOstar Omega,BMG Labtech, Germany) every 6 minutes during 72 hours, at 22°C under shaking at 700 rpm using a linear shaking mode. Three to six independent biological repeats were performed for each strain.

### Host genetic variation in response to bacterial isolates

To estimate the level of genetic variation of *A. thaliana* in response to each of the 22 selected strains in *in vitro* conditions, seven accessions were chosen to represent the genetic and habitat diversity observed among the 163 natural populations (Frachon et al., 2018, 2019) (Supplementary Figure 2). The reference accession Col-0 was also included in the *in vitro* experiments. To test whether the level of genetic variation of *A. thaliana* in response to OTUs was dependent on the developmental stage at which plants were inoculated, the eight accessions were inoculated at the seed and seedling stages.

#### Seed sterilization

Seeds were surface-sterilized with chlorine gas one day before sowing and kept at 4°C. For this, eight Ø5cm Petri dishes, each containing seeds from one accession, and a beaker with 50 mL of bleach were placed in a plastic box under a fume hood. A volume of 5 mL of 37% hydrochloric acid was added to the bleach before closing the plastic box. After three hours, Petri dishes were opened in a microbiological sterile fume hood and left two hours to allow the evaporation of the bleach steam.

#### Plant growth conditions

Plants were grown in 48-well and 24-well plates for the inoculation at the seed and seedling stages, respectively, on a 0.5x MS medium (Murashige & Skoog medium) modified according to Dr Andy Gloss (Department of Biology, New-York University). This medium contains 2.2 g of MS medium, 0.5 g of 2-(N-Morpholino)-ethanesulfonic acid, 6.0 g of plant tissue culture agar (PhytoTechnology Laboratories, A296), 1L of milli-Q water and a pH adjusted to range of 5.7-5.8. A volume of 700 µL and 1 mL was dispensed in each well of the 48-well and 24-well plates, respectively. A single seed was sown in each well. Seeds were stratified at 4 °C in the dark for seven days. Plates were then placed in a phytotron with short day conditions (10H photoperiod, light intensity ∼ 80 µmol m^-2^ s^-1^, 21°C, 50% hygrometry). To reduce the effects of micro-environmental variation, the position of the plates was randomized in the phytotron every other day of the experiment.

#### Experimental design

For the inoculation at the seed stage, the eight accessions of *A. thaliana* were inoculated with the 22 selected strains, according to a split-plot design with two blocks. Each block contained 24 48-well plates, corresponding to two plates for the mock inoculation and one plate for each of the 22 strains. For each ‘block*treatment’ combination, six seeds of the eight accessions were randomly sown among the 48 wells.

For the inoculation at the seedling stage, the eight accessions of *A. thaliana* were inoculated with the 22 selected strains, according to a split-plot design with four blocks. Each block contained 24 24-well plates, corresponding to two plates for the mock inoculation and one plate for each of the 22 strains. For each ‘block*treatment’ combination, three seeds of the eight accessions were randomly sown among the 24 wells.

In both experiments, randomization was kept identical among treatments within a block, but differed between blocks.

#### Inoculation

Bacterial suspensions were prepared as for the growth kinetics but adjusted to an OD600nm of 0.5 in sterile water and 0.2 in 0.001 % (v/v) of Tween 20, for the seed and seedling inoculation, respectively. For inoculation at the seed stage, 5 µL of inoculum were dropped off on each seed the day of the sowing. For inoculation at the seedling stage, seedlings were inoculated 18 days after sowing using a ‘droplet’ method, which consisted of dispensing 50 µL on each rosette. Sterile water without or with 0.001 % (v/v) of Tween 20 was used for the mock corresponding treatments. Plates were closed with a micropore tape (Micropore Surgical tape 3M 1530-0) to limit water evaporation of the medium, and incubated in the same phytotron conditions as described above.

#### Phenotyping

For the two experiments, the germination date was scored on a daily basis between three and seven days after sowing. For inoculation at the seed stage, the 48-well plates were scanned at 14, 21 and 28 days after inoculation (dai) using the scanner Object Scan 1600 scanner (Microtek® Taiwan). For inoculation at the seedling stage, 24-well plates were scanned one day before inoculation (dbi) and 7, 14 and 21 dai using the same scanner. Based on the pictures, the vegetative growth of each plant was manually scored using a scale ranging from one (very small plants) to seven and eight (well-grown plants) for the inoculation at the seed and seedling stages, respectively. For each experiment, a total of 2,304 plants were phenotyped after inoculation. No disease symptoms was observed in our growing conditions.

#### Statistical analysis

For both experiments, we explored the genetic variation among the eight accessions of *A. thaliana* in response to each of the 22 bacterial strains, using the following mixed model (PROC MIXED procedure in SAS v. 9.4, SAS Institute Inc., Cary, NC, USA):

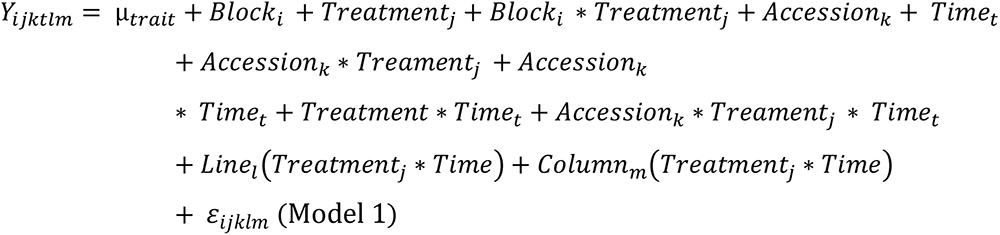

where Y corresponds to the score of plant development, ‘µ’ is the overall mean of the phenotypic data, ‘Block’ accounts for differences in micro-environmental conditions between the blocks, ‘Line(Treatment)’ and ‘Column(Treatment)’ accounts for difference in micro-environmental conditions within 48-well or 24-well plates, ‘Treatment’ corresponds to the effect of each bacterial strain in comparison with the mock treatment, ‘Accession’ corresponds to the genetic differences among the eight accessions, ‘Time’ expressed in dbi/dai accounts for the evolution of developmental stage along the course of the experiment, the interacting terms test whether pairwise genetic differences among the eight accessions differ between the mock treatment and each treatment with a bacterial strain and/or with time, and ‘ε’ is the residual term.

All factors were treated as fixed effects. For calculating *F* values, terms were tested over their appropriate denominators. Given the split-plot design used in the two experiments, the variance associated with the ‘block * treatment’ term was for example used as the error term for testing the ‘block’ and ‘treatment’ effects. A correction for the number of tests was performed to control the False Discover Rate (FDR) at a nominal level of 5%.

For both *in vitro* experiments, least-square means (LSmeans) of the eight accessions were estimated for each ‘treatment * scoring time’ combination using the following model (PROC MIXED procedure in SAS v. 9.4, SAS Institute Inc., Cary, NC, USA):

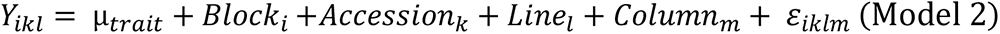

Because *A. thaliana* is a highly selfing species (Platt et al., 2010), LSmeans correspond to genotypic values of accessions.

## RESULTS

### Isolation of strains of the most prevalent and abundant OTUs of the leaf compartment in 163 natural populations of *A. thaliana* located south-west of France

Based on the characterization of bacterial communities of 821 rosettes of *A. thaliana* collected *in situ* in autumn 2014 and spring 2015 from 163 local populations located south-west of France (Bartoli et al., 2018), we focused on 12 non-pathogenic OTUs displaying both high prevalence (ranging from 5.7% for OTU2 to 44.6% for OTU1, mean = 19.1%) and high mean relative abundance (MRA) (ranging from 0.7% for OTU12 to 16.1% for OTU1, mean = 3.1%) (Table 1). Based on *gyrB* taxonomic affiliation, these 12 OTUs belongs to nine genera, *i.e. Acidovorax*, *Caulobacter*, *Collimonas, Methylobacterium, Modestobacter, Paraburkholderia*, *Pseudomonas*, *Sphingomonas* and *Variovorax* (Table 1, Supplementary Data Set 1). The 12 candidate OTUs are well spread across the *gyrB*-related phylogeny of the 30 most abundant OTUs across the 163 local populations of *A. thaliana* by considering both leaf and root compartments (Bartoli et al., 2018) (Figure 1A).

To isolate representative strains for each of the 12 candidate OTUs, we adopted two approaches (Supplementary Figure 1). Firstly, based on a community-based culture (CBC) approach, we isolated 7,259 bacterial colonies from 152 natural populations of *A. thaliana*. Using the Illumina technology, the amplification of a *gyrB* amplicon was successfully achieved for 7,066 bacterial colonies, from which 3,746 (*i.e.* 53%) were taxonomically assigned at the phylum level. Among these 3,746 samples, we identified five samples with at least 10% of the reads belonging to an ASV matching to OTU2 with an identity ≥ 98%. With the same criteria, we identified 55 samples for OTU3, 21 samples for OTU5, 365 samples for OTU6 and 59 samples for OTU29. To isolate pure strains from these samples and pure strains for the remaining seven OTUs, we adopted a second approach based on selective media, antibiotic resistance and the design of specific primers (Supplementary Figure 1). This informative-driven approach allowed the isolation of 35 pure strains for seven OTUs (3 strains for OTU2, 7 strains for OTU3, 1 strain for OTU4, 8 strains for OTU5, 6 strains for OTU6, 2 strains for OTU13 and 8 strains for OTU29). These seven OTUs show diverse morphological aspects in terms of color and shape (Figure 1B).

### Genomic variation among and within OTUs

As a first step to characterize genetic variation among and within OTUs, we generated *de novo* genome sequences using PacBio technology for 22 strains maximizing the geographic diversity for each OTU (3 strains for OTU2, four strains for OTU3, 1 strain for OTU4, 3 strains for OTU5, 6 strains for OTU6, 2 strains for OTU13 and 3 strains for OTU29). Despite several attempts, we failed to generate *de novo* genome sequence for one strain of OTU3 and one strain of OTU13. For all the other strains, chromosome and plasmid genomes were assembled in single contigs and circularized, with the exception of one plasmid for the strain of OTU13 that was not circularized (Table 2). Strong genomic differences were observed among the seven OTUs (Table 2). Firstly, the bacterial chromosome size varies from ∼3.2Mb for OTU4 to ∼7.7Mb for one strain of OTU3 (mean = 5.5Mb). Secondly, because the number and size of plasmids largely differed among the seven OTUs and even among strains within some OTUs (Table 2), the total genome size was not strictly related to the chromosome size. For instance, the total size of plasmids represents on average 46.8% and 29.3% of total genome size for OTUs 2 and 4, respectively. On the other hand, for the OTUs 6, 13 and 29 with the presence of plasmids in at least one strain, the total size of plasmids represents less than 5% of the total genome size. Thirdly, G+C content varies between 60% for OTUs 5 and 6 to 70% for OTU4 (mean = 64.4%). Fourthly, the number of genes largely differs among OTUs (from ∼3,723 genes for OTU29 to ∼8,426 genes for OTU2) and was strictly correlated with total genome size (Pearson’s correlation coefficient *r* = 0.99). Finally, the percentage of coding sequences with assigned function varies between 54.3% for OTU13 to 67.9% for OTU4 (mean = 64.3%).

**Table 2.** Summary of the assembly and annotation characteristics of the 20 whole-genome sequenced strains belonging to eight OTUs.

| Strains | Total assembly size (bp) | Number of contigs | Number of chromosomes | Chromosome size (Mb) | Number of plasmids | Plasmids size (Kb) | G+C content (%) | Number of genes | Number of CDSs (coding) | rRNA | tRNA | Number of CDSs with assigned function | Number of CDSs without assigned function |
| --- | --- | --- | --- | --- | --- | --- | --- | --- | --- | --- | --- | --- | --- |
| OTU2_*Pfu*_1 | 9,357,004 | 6 | 2 | 4.78 + 3.53 | 4 | 435 + 295 + 228 + 84 | 62% | 8624 | 8465 | 12 | 74 | 5384 (63.6%) | 3081 (36.4%) |
| OTU2_*Pfu*_2 | 9,120,800 | 4 | 2 | 4.94 + 3.58 | 2 | 447 + 147 | 62% | 8329 | 8171 | 12 | 71 | 5313 (65.0%) | 2858 (35.0%) |
| OTU2_*Pfu*_3 | 9,122,278 | 4 | 2 | 4.94 + 3.59 | 2 | 447 + 147 | 62% | 8327 | 8169 | 12 | 71 | 5313 (65.04%) | 2856 (34.96%) |
| OTU3a_*Oxa*_1 | 7,481,272 | 1 | 1 | 7.48 | 0 |  | 64% | 6337 | 6208 | 14 | 86 | 3969 (63.9%) | 2239 (36.0%) |
| OTU3a_*Oxa*_2 | 7,693,415 | 1 | 1 | 7.69 | 0 |  | 64% | 6514 | 6382 | 14 | 89 | 4055 (63.5%) | 2327 (36.46%) |
| OTU3b_*Oxa*_1 | 6,868,374 | 1 | 1 | 6.87 | 0 |  | 64% | 5935 | 5795 | 14 | 97 | 3657 (63.1%) | 2138 (36.9%) |
| OTU4_*Com*_1 | 4,655,332 | 5 | 1 | 3.29 | 4 | 493 + 424 + 290 + 159 | 70% | 4268 | 4183 | 6 | 53 | 2842 (67.9%) | 1341 (32.1%) |
| OTU5_*Pmo*_1 | 6,004,654 | 1 | 1 | 6.00 | 0 |  | 60% | 5446 | 5272 | 16 | 72 | 3526 (66.9%) | 1746 (33.1%) |
| OTU5_*Pmo*_2 | 5,969,148 | 1 | 1 | 5.97 | 0 |  | 60% | 5412 | 5238 | 16 | 71 | 3502 (66.9%) | 1736 (33.1%) |
| OTU5_*Pmo*_3 | 5,980,820 | 1 | 1 | 5.98 | 0 |  | 60% | 5416 | 5242 | 16 | 70 | 3528 (67.3%) | 1714 (32.7%) |
| OTU6_*Psi*_1 | 5,988,833 | 1 | 1 | 5.99 | 0 |  | 60% | 5458 | 5282 | 16 | 73 | 3539 (67.0%) | 1743 (33.0%) |
| OTU6_*Psi*_2 | 6,005,098 | 1 | 1 | 6.00 | 0 |  | 60% | 5401 | 5227 | 16 | 72 | 3520 (67.5%) | 1697 (32.5%) |
| OTU6_*Psi*_3 | 5,986,784 | 1 | 1 | 5.99 | 0 |  | 60% | 5431 | 5258 | 16 | 71 | 3522 (67.0%) | 1736 (33.0%) |
| OTU6_*Psi*_4 | 5,915,411 | 1 | 1 | 5.91 | 0 |  | 60% | 5377 | 5205 | 16 | 73 | 3513 (67.5%) | 1692 (32.5%) |
| OTU6_*Psi*_5 | 6,095,266 | 2 | 1 | 5.99 | 1 | 103 | 60% | 5597 | 5422 | 16 | 74 | 3562 (65.7%) | 1860 (34.3%) |
| OTU6_*Psi*_6 | 5,918,345 | 1 | 1 | 5.92 | 0 |  | 60% | 5423 | 5247 | 16 | 75 | 3510 (66.9%) | 1737 (33.1%) |
| OTU13_*Msp*_1 | 5,074,646 | 5 | 1 | 4.90 | 4 | 56 + 42 + 40 + 38 | 69% | 4903 | 4800 | 15 | 64 | 2608 (54.3%) | 2192 (45.7%) |
| OTU29_*Sph*_1 | 3,810,917 | 3 | 1 | 3.62 | 2 | 126 + 65 | 66% | 3629 | 3551 | 9 | 55 | 2103 (59.2%) | 1448 (40.8%) |
| OTU29_*Sph*_2 | 3,825,112 | 2 | 1 | 3.76 | 1 | 65 | 66% | 3610 | 3532 | 9 | 56 | 2128 (60.2%) | 1404 (39.8%) |
| OTU29_*Sph*_3 | 4,127,732 | 2 | 1 | 4.00 | 1 | 74 | 66% | 3931 | 3851 | 9 | 58 | 2258 (58.6%) | 1593 (41.4%) |

ANI-based genomic taxonomic classification confirmed or refined the *gyrB*-based taxonomic classification for OTU2, OTU5, OTU6 and OTU13 as *Paraburkholderia fungorum*, *Pseudomonas moraviensis*, *Pseudomonas siliginis* and *Methylobacterium* sp., respectively (Table 1, Supplementary Data Set 6). However, ANI-based genomic taxonomic classification for the remaining three OTUs only provided a resolution down to the family level, *i.e.* Oxalobacteraceae for OTU3, Comamonadaceae for OTU4 and Sphingomonadaceae for OTU29 (Table 1, Supplementary Data Set 6). For each OTU, pairwise ANI values were computed between all strains and revealed that all strains belong to the same species (using a species cut-off value of 0.95), with the exception of OTU3 for which two species were identified (Supplementary Table 4). Hereafter, these two species are named OTU3a_*Oxalobacteraceae* bacterium and OTU3b_*Oxalobacteraceae* bacterium (Table 1), thereby leading to a total number of eight OTUs.

A strong variation in the gene content was also observed among strains within a given OTU. For instance, 5,185 orthogroups were identified among the six strains of OTU6_*Pseudomonas siliginis*, with 83.1% (4,310 orthogroups) being shared among the six strains and 8.8% (456 orthogroups) being specific to one or two strains (Figure 2).

**Figure 2.**
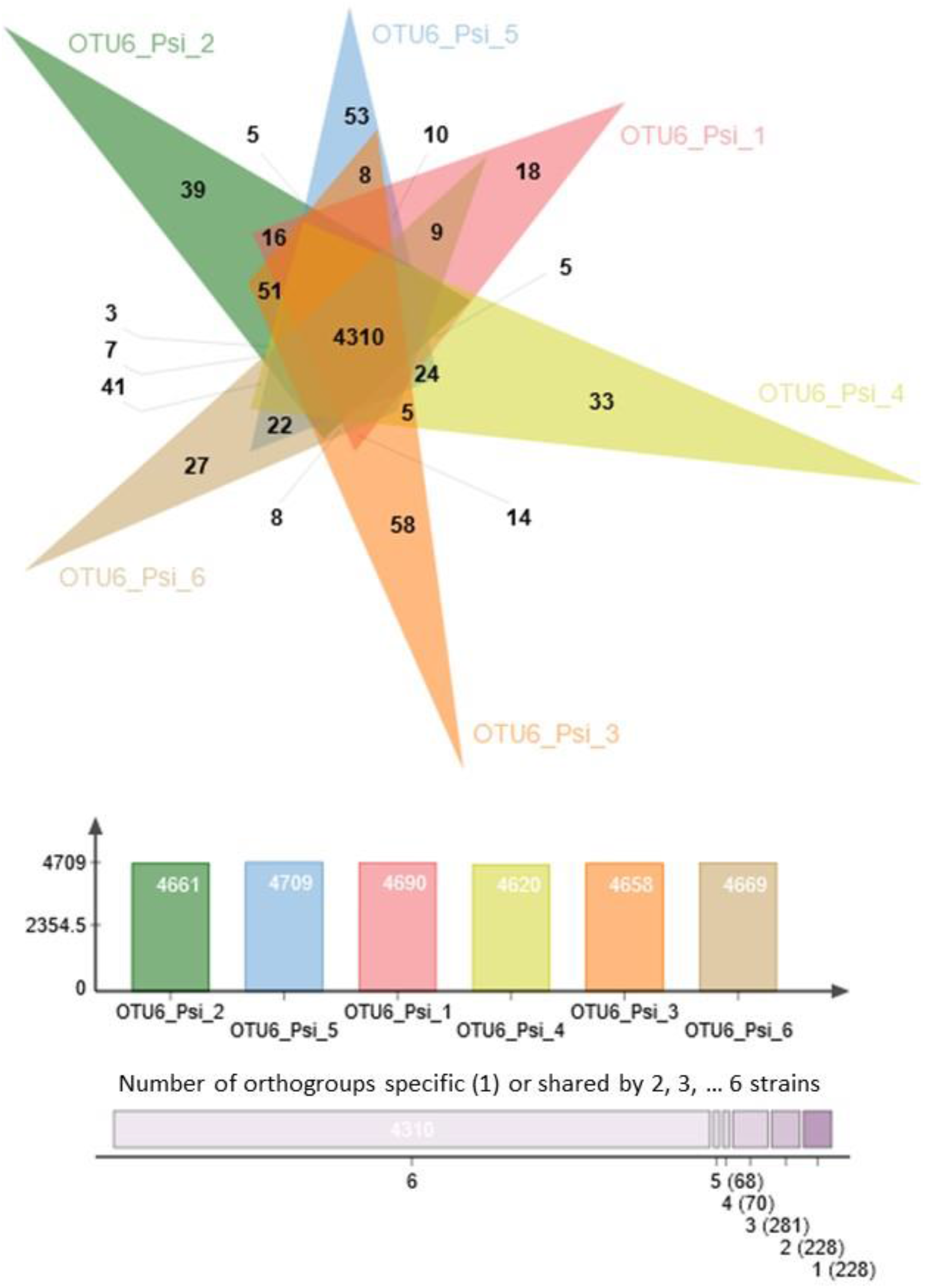
Venn diagram illustrating the number of common and specific orthogroups among the six strains of OTU6_*Pseudomonas siliginis*. Colored bars indicate the number of orthogroups for each strain. The vertical stacked bar plot indicates the number of orthogroups specific to one strain (right) or shared between two to six strains.

### Inter- and intra-genetic variation of bacterial growth kinetics

To further characterize our set of 22 representative strains, we monitored their growth kinetics on two media with contrasting nutrient availability, *i.e.* the R2A minimal medium and the TSB rich medium. Over the time course of 40 hours, growth kinetics was more diverse among the eight OTUs in the rich medium than in the minimal medium (Figure 3). In the rich medium, on average, the OTUs 2, 5 and 6 reached larger population size than the other OTUs, the two strains of OTU13 growing very slowly (Figure 3B). The level of genetic variation among strains within a specific OTU for growth kinetics was highly dependent on the ‘medium * OTU identity’ combination. While almost no intra-OTU genetic variation was detected in both media for OTU6 (Figure 3) and OTU5 (Supplementary Figure 3), intra-OTU genetic variation was detected in both media for OTUs 3a, 3b (Figure 3) and OTU13 (Supplementary Figure 3). For the OTUs 2 and 29, the level of genetic variation among strains was highly dependent on the medium. For instance, while the three strains of OTU29 had a similar growth kinetics on the minimal medium, they largely differed in their growth kinetics on the rich medium starting after ∼15H of culture (Figure 3). An opposite pattern was observed for OTU2, with genetic variation for growth kinetics observed in the minimal medium but not in the rich medium (Supplementary Figure 3).

**Figure 3.**
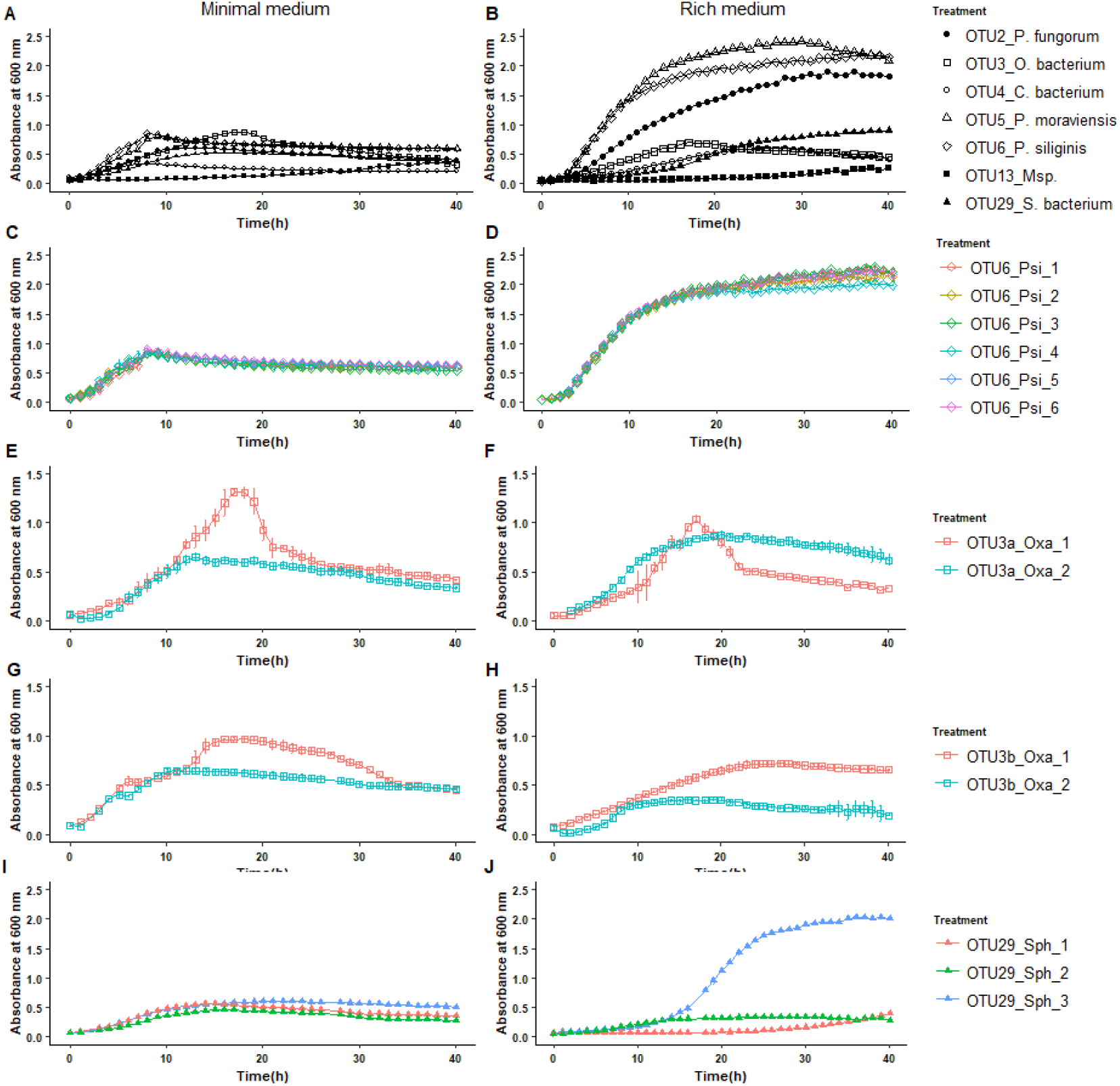
Inter- and intra-OTU genetic variation for growth kinetics on two media with contrasting nutrient availability. **(A)** Variation among the OTUs for growth kinetics on the R2A minimal medium. **(B)** Variation among the OTUs for growth kinetics on the TSB rich medium. **(C)**, **(E)**, **(G)** and **(I)** Intra-OTU genetic variation for the OTUs 6, 3a, 3b and 29 on the R2A minimal medium. **(D)**, **(F)**, **(H)** and **(J)** Intra-OTU genetic variation for the OTUs 6, 3a, 3b and 29 on the TSB rich medium.

### The extent of host genetic variation in response to bacterial strains in *in vitro* conditions

To estimate the extent of GxG interactions between *A. thaliana* and the main members of its microbiota, seven local accessions of *A. thaliana* originating from diverse habitats across the 163 natural populations located south-west of France (Supplementary Figure 2) as well as the reference accession Col-0, were inoculated with the 22 selected bacterial strains at the seed and seedling stages in *in vitro* conditions (Figure 4A, Figure 5A).

**Figure 4.**
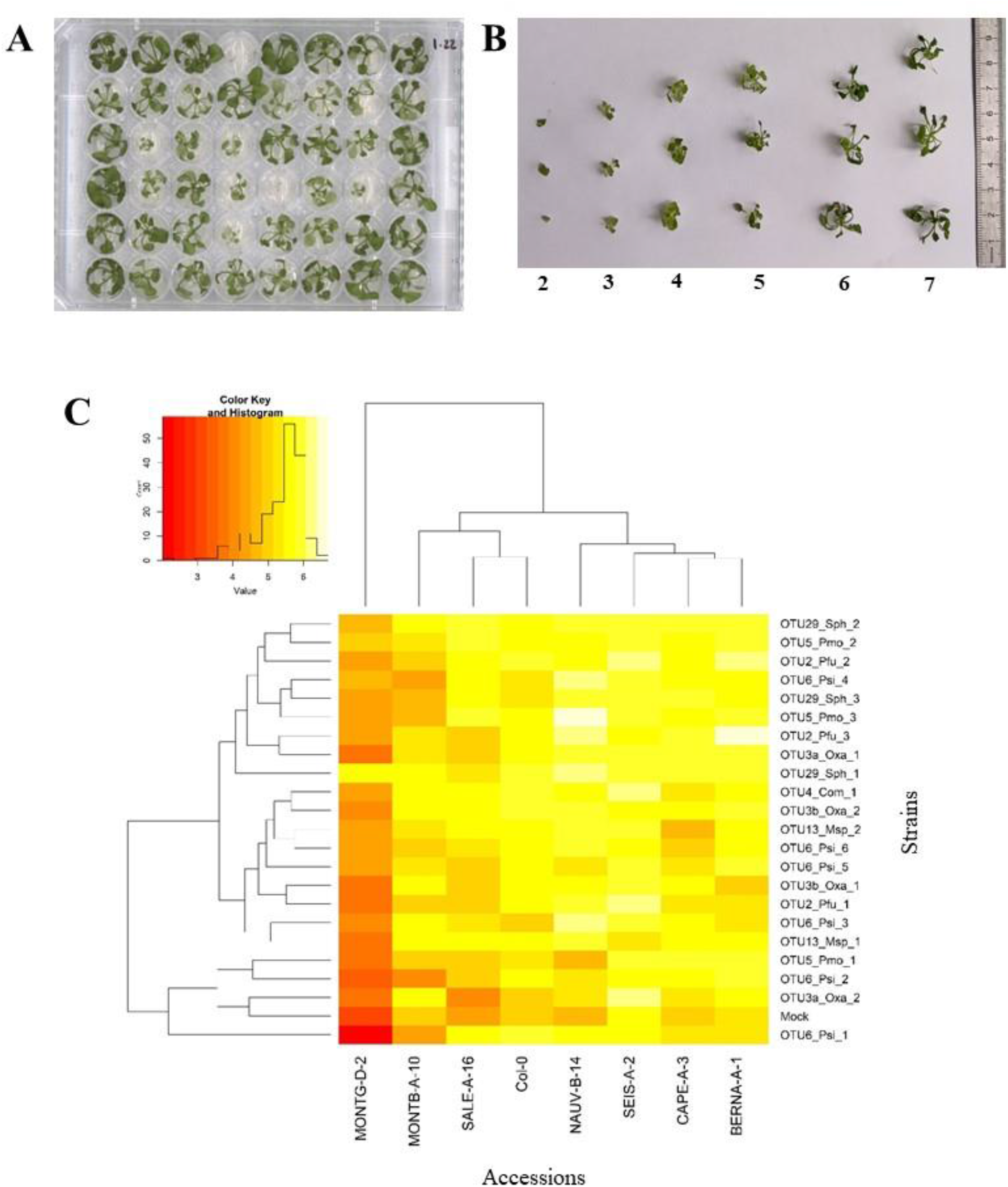
Genetic variation among eight *A. thaliana* accessions in response to 22 bacterial strains inoculated at the seed stage in *in vitro* conditions. **(A)** Photography illustrating plants grown in 48-well plates and scored *in vitro* at 28 dai. **(B)** Scoring scale of plant development after inoculation at the seed stage, ranging from 1 (very small and/or not healthy plant) to 7 (very well-grown and/or healthy plant). **(C)** Double hierarchical clustering based on genotypic values, illustrating the genetic variation among the eight *A. thaliana* accessions in response to the 22 representative bacterial strains belonging to eight OTUs 28 dai. Inner plot: Histogram illustrating the distribution of genotypic values for plant development according a scale from 1 to 7.

**Figure 5.**
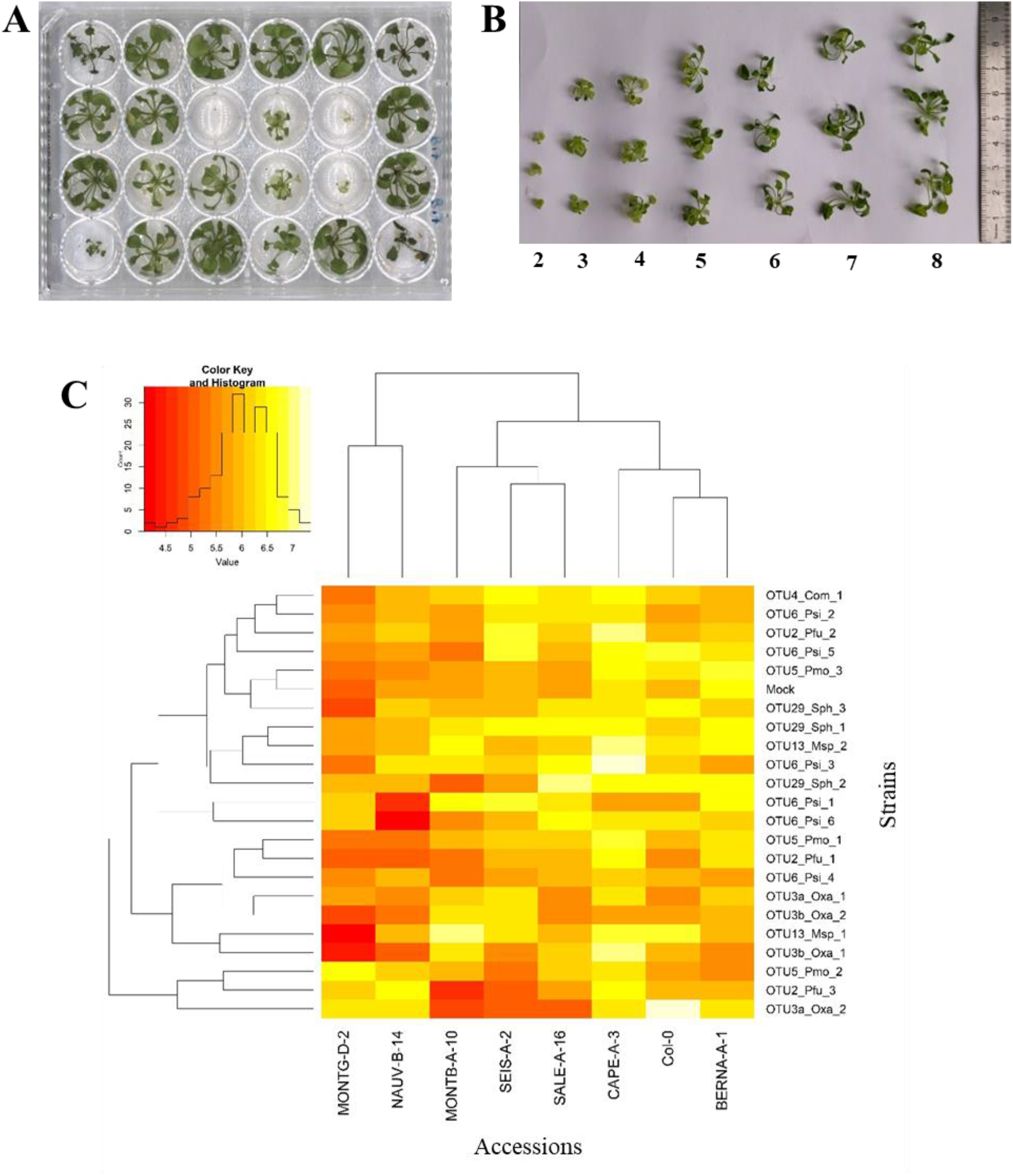
Genetic variation among eight *A. thaliana* accessions in response to 22 bacterial strains inoculated at the seedling stage in *in vitro* conditions. **(A)** Photography illustrating plants grown in 24-well plates and scored *in vitro* at 21 dai. **(B)** Scoring scale of plant development after inoculation at the seed stage, ranging from 1 (very small and/or not healthy plant) to 8 (very well-grown and/or healthy plant). **(C)** Double hierarchical clustering based on genotypic values illustrating the genetic variation among the eight *A. thaliana* accessions in response to the 22 representative bacterial strains belonging to eight OTUs 21 dai. Inner plot: Histogram illustrating the distribution of genotypic values for plant development according a scale from 4 to 7.5.

Based on scoring vegetative growth following previously defined scales (Figure 4B, Figure 5B), several patterns emerged from these *in vitro* experiments. Firstly, when significant, host genetic variation in response to a particular bacterial strain, was not dependent on the timing of scoring (Supplementary Tables 5 & 6). Secondly, host genetic variation was dependent on the developmental stage at which plants were inoculated (Figure 4C, Figure 5C, Supplementary Figure 4). No significant variation among the eight accessions of *A. thaliana* was detected in response to 10 bacterial strains, when inoculated either at the seed stage or at the seedling stage (Supplementary Tables 5 & 6). For the remaining 12 bacterial strains, significant genetic variation among the eight accessions was detected only when inoculated at the seed stage (n = 4 strains), only when inoculated at the seedling stage (n = 5 strains), or when inoculated at both stages (n = 3 strains) (Supplementary Table 5 and 6). Thirdly, host genetic variation was more dependent on the strain identity than the OTU identity (Figure 4C, Figure 5C, Supplementary Figure 4). For instance, while no host genetic variation was detected in response to OTU2_*Pfu*_1 and OTU2_*Pfu*_2, genetic variation among the eight accessions was detected in response to OTU2_*Pfu*_3 at both developmental stages (Supplementary Tables 5 and 6). Another example is related to OTU29_*Sphingomonadaceae*_bacterium. When inoculated at the seed stage, host genetic variation was observed in response to OTU29_*Sph*_1 and OTU29_*Sph*_3 but not in response to OTU29_*Sph*_2 (Supplementary Table 5). An opposite pattern was observed when inoculation was conducted at the seedling stage (Supplementary Table 6). Fourthly, contrasted responses were observed among the eight accessions in response to a particular bacterial strain (Figure 4C, Figure 5C, Supplementary Figure 4). For instance, at 28 dai after seed inoculation, the strain OTU6_*Psi*_1 had a negative effect on the accessions MONTG-D-2 and MONTB-A-10, a neutral effect on the accessions BERNA-A-1 and SEIS-A-2, and a positive effect on the four remaining accessions (Figure 4C). Similarly, at 21 dai after seedling inoculation, the strain OTU3a_*Oxa*_2 had a positive effect on the accessions MONTG-D-2, NAUV-B-14 and Col-0, a neutral effect on the accession CAPE-A-3, and a negative effect on the four remaining accessions (Figure 5C).

## DISCUSSION

### Mining ecologically relevant genetic diversity within leaf-associated bacterial OTUs of *A. thaliana*

The importance of studying the functionality of microbiota and establishing causal relationships between microbial species and between a given host plant and its microbiota mainly relies on adopting reductionist approaches (Vorholt et al., 2017; Hassani et al., 2018; Liu et al., 2019; Fitzpatrick et al., 2020), which first requires the establishment of representative microbial collections. In *A. thaliana*, such collections of bacterial and/or fungal isolates have been generated in several instances (Bai et al., 2015; Lebeis et al., 2015; Qi et al., 2021), which in turn allowed establishing SynComs (e.g. Durán et al., 2018; Carlström et al., 2019). While informative, these microbial collections would certainly benefit from exploiting intra-OTU genetic diversity from a large range of environments inhabited by the host species in order to increase the ecological realism of SynComs, as previously suggested for pathogens (Bartoli et al., 2016).

In this study, we adopted two complementary approaches to isolate pure strains from the 12 most abundant and prevalent non-pathogenic leaf-associated bacterial OTUs in *A. thaliana* located south-west of France. Using the CBC approach, recovery estimates of the most abundant 6,627 OTUs identified among the 163 natural populations of *A. thaliana* (Bartoli et al., 2018) were 53%, a value similar to the recovery estimate (54%) of the top 100 leaf-associated bacterial OTUs from *A. thaliana* plants collected from natural sites in France, Germany and Switzerland (Bai et al., 2015). However, by using the CBC approach, we failed to isolate colonies for seven out of our 12 most abundant and prevalent leaf-associated bacterial OTUs. Because we obtained Illumina reads for 97.3% of the 7,259 picked colonies, this failure seems not to result from our colony PCR based protocol. However, it may originate from (i) the use of only one medium (TSB) for the CBC approach, and (ii) the use of leaf samples for strain isolation that differ from the leaf samples used for characterizing bacterial communities based on a *gyrB*-based metabarcoding approach. In addition, despite a large number of growing conditions implemented in this study coupled with the design of specific primers (Supplementary Table 1), our informative-driven approach only allowed isolating strains from two additional OTUs. Our capacity of isolate strains for only seven out of our 12 candidate OTUs provides another example of the difficulty of establishing a representative collection of microbes. The un-cultivability of many microbes could result from the absence of specific nutrients in the medium, specific factors produced by other microbes that are necessary for the OTU of interest (*i.e.* cross-feeding), rigorous negative interspecies interactions (such as competition and inhibition), slow growth and dormancy (Nemr et al., 2021).

Optimizing the processes of isolating microbial strains can rely on the use of plant extract as medium, thereby providing specific nutrients for the OTUs of interest. For instance, the use of plant extract added to the medium led to a slight increase in microbial diversity and a sharp increase of viable bacteria in both minimal and rich media (Eevers et al., 2015). Such an approach has optimized the cultivability of rhizobacteria and supported recovery from plant-soil environments (Youssef et al., 2016). Adding to the medium plant extracts of *A. thaliana* accessions from the natural populations in which the OTU of interest is largely present, may help to isolate strains from our five remaining strain-free leaf-bacterial associated OTUs. In this context, it is urgent to develop novel culturing methods mimicking the host habitat. Few examples have been reported with the isolation chips (Berdy et al., 2017) developed on soil and sediments, but more efforts are still needed on the plant leaf compartment. Culturomics, the high-throughput isolation coupled to sequencing and mass-spectrometry, will definitely help in improving bacterial collections associated with phyllosphere.

### Extensive genetic and genomic diversities within leaf-associated bacterial OTUs

Extensive genetic and genomic diversity was previously described among strains of the main pathogenic bacterial species collected in natural populations of *A. thaliana*, including *P. syringae* and *P. viridiflava* (Karasov et al., 2014; Bartoli et al., 2018; Karasov et al., 2018), *X. arboricola* (Wang et al., 2018) and *X. campestris* (Bartoli et al., 2018). In this study, we also detected an extensive genetic diversity among strains within the leaf-associated bacterial OTUs considered in this study, at the genome level, for their growth capacities and for their host responses. This highlights the importance of investigating within-OTU genetic diversity in order to better understand the functionality of bacteria-bacteria interactions as well as the interplay between *A. thaliana* and members of its bacterial communities.

Genome sequencing confirmed or refined the *gyrB*-based taxonomic affiliation of four OTUs, *i.e. P. fungorum* (OTU2), *Methylobacterium* sp. (OTU13) and the two *Pseudomonas* species *P. moraviensis* (OTU5) and *P. siliginis* (OTU6). All these four bacterial species have been shown to act as biocontrol agents, to affect root development, to promote vegetative growth and ultimately yield, of diverse plants such as *A. thaliana*, potato, strawberry, tomato ad wheat (Hultberg et al., 2010; Ul Hassan and Bano, 2015; Rafikova et al., 2016; Klikno and Kutschera, 2017; Rahman et al., 2018; Grossi et al., 2020). In addition, both *P. moraviensis* and *P. siliginis* have been identified as the main candidate bacterial species controlling most members of the root and leaf bacterial pathobiota, in particular *P. viridiflava* and *X. campestris*, across natural populations of *A. thaliana* located south-west of France (Bartoli et al., 2018). On the other hand, despite a deeper taxonomic resolution provided by the *gyrB* gene rather than the *16S rRNA* gene (Barret et al., 2015; Rezki et al., 2016), the *gyrB*-based taxonomic affiliation of OTU3, OTU4 and OTU29 was only confirmed at the family level by genome sequencing. In addition, genome sequencing revealed the presence of two bacterial species in OTU3. Altogether, these results suggest the identification of four novel leaf-associated bacterial species in *A. thaliana*. Confirming these novel bacterial species would require chemotaxonomic methods, albeit recent studies argue that a genome sequence suffices to prove that a strain represents a novel species or not (Vandamme and Sutcliffe, 2021).

Similarly to bacterial pathogens, sequencing several strains from the same non-pathogenic bacterial species revealed a non-negligible fraction of genes that are not common to all strains, as exemplified for *P. siliginis*. For a given non-pathogenic bacterial species, sequencing the genome of several tens additional strains will reveal the relative fraction of the core and accessory genomes. This in turn may help to identify the conserved metabolic pathways necessary for bacterial growth in any environment and the genes involved in specific interactions with its abiotic and biotic environment, respectively.

In agreement with the genomic variation observed within the OTUs, we detected genetic variation among strains for host response, which was itself both dependent on the identity of the *A. thaliana* accession and the plant developmental stage at which strains were inoculated. This pattern may result from the construction of contrasted ecological niches between accessions of *A. thaliana* that additionally differ between the seed and seedling stage, thereby affecting the spectrum of nutrients available for the growth of a specific strain. This hypothesis is reinforced by the genetic variation of growth kinetics observed among strains either in one or both of the two media tested. Assessing our 22 candidate bacterial strains for both their diversity of substrate utilization and their growth kinetics on plant extracts harvested on diverse *A. thaliana* accessions at complementary developmental stages, may help to better understand the interplay between within-OTU genetic variation and nutrient availability conditioning on host genetics and development.

Interestingly, for most strains tested in *in vitro* conditions, both positive and negative growth responses were observed between the eight accessions of *A. thaliana* for a specific strain, whatever the plant developmental stage considered. In our study, declaring a strain as a plant growth-promoting bacteria (PGPB) would have been highly dependent on the host genetics. On the other hand, the large genetic variation observed within *A. thaliana* in response to most bacterial strains provides a unique opportunity to set up Genome-Wide Association studies based on the 500 whole-genome sequenced accessions located south-west of France (Frachon et al., 2018), in order to decipher the genetic and molecular mechanisms involved in the ecologically relevant dialog between *A. thaliana* and the main members of its leaf microbiota.

## DATA AVAILABILITY STATEMENT

The whole genome sequences of the 20 bacterial strains have been deposited at DDBJ/ENA/GenBank. The BioProject accession number is PRJNA848627. The raw phenotypic data supporting the conclusions of this article will be made available by the authors, without undue reservation.

## ACKNOWLEDGMENTS

D.R.S. was funded by a PhD fellowship from CONACyT. R.D. was funded by a grant from the French Ministry of National Education and Research. We are grateful to Carine Huard-Chauveau and Tatiana Vernié for their help with isolating the 7,259 bacterial colonies, and to Andy Gloss for advices on plant culture methods. We thank Cécile Pouzet of the TRI-FRAIB Imaging platform facilities, FR AIB 3450, CNRS-UTIII, for advices for scan measurements. This study was performed at the LIPME belonging to the Laboratoire d’Excellence (LABEX) entitled TULIP (ANR-10-LABX-41).

## AUTHOR CONTRIBUTIONS

Conceptualization, F.V., F.R., D.R.-S.; Methodology, D.R.-S., C.G., B.M., E.B., F.V. and F.R.; Formal Analysis, D.R.-S., C.G., B.M., R.D., V.P., S.C, F.V. and F.R.; Investigation, D.R.-S., C.G., B.M., R.D., E.B., FV. And F.R.; Resources, C.B., F.R., and B.M.; Data Curation: V.P., S.C. and C.B.; Writing – Original Draft, D.R.-S., C.G., B.M., S.C., F.V. and F.R.; Writing – Review & Editing, D.R.-S., C.G., B.M., R.D., C.B., F.V. and F.R. Funding Acquisition, D.R.-S., F.V. and F.R.; Supervision, F.V. and F.R.

## CONTRIBUTION TO THE FIELD STATEMENT

The potential of harnessing the microbiome towards the improvement of plant health is an ever-growing interest to propose innovative, sustainable and eco-friendly agricultural systems. While an uncountable number of studies on taxonomic profiling of microbial communities revealed key patterns of microbiota assemblages in plants, both studying the functionality of microbiota and establishing causal relationships between plants and non-pathogenic microbes remains a lofty challenge, which is primarily based on representative collections of microbes isolated from the diversity of environments inhabited by the plant of interest. Most microbial collections from *A. thaliana* were built on the isolation of one strain per OTU from a limited number of sites. Neglecting the ecological potential of genetic diversity within microbial species may strongly impact the outcomes of plant-microbiota interactions In this study, we revealed extensive genetic variation between strains within the most abundant and prevalent non-pathogenic leaf-associated bacterial species in French natural populations of *A. thaliana*, at the genomic level, for their growth capacity and their host response. Altogether, our results highlight the need to consider genetic variation within non-pathogenic bacterial species for a better understanding of plant-microbiota interactions and the underlying genetic and molecular mechanisms.

## Supplementary Material including four Supplementary Figures and four Supplementary Tables

**Supplementary Figure 1** | CBC and informative-driven approaches to isolate representative strains of the 12 most abundant and prevalent leaf OTUs across 163 natural populations of *Arabidopsis thaliana* located south-west of France.

**Supplementary Figure 2** | Pictures illustrating the habitats of the seven accessions chosen to test host genetic variation in response to bacterial isolates.

**Supplementary Figure 3** | Intra-OTU genetic variation for growth kinetics on two media with contrasting nutrient availability. **(A)**, **(C)**, **(E)** and **(G)** Intra-OTU genetic variation for the OTUs 4, 5, 13 and 2 on the R2A minimal medium. **(B)**, **(D)**, **(F)** and **(H)** Within-OTU genetic variation for the OTUs 4, 5, 13 and 2 on the TSB rich medium.

**Supplementary Figure 4** | Double hierarchical clustering based on genotypic values, illustrating the genetic variation among the eight *A. thaliana* accessions in response to the 22 representative bacterial strains belonging to eight OTUs at 28 dai. **(A**) Inoculation at the seed stage with scoring at 14 dai (top panel) and 21 dai (bottom panel). **(B**) Inoculation at the seedling stage with scoring at 7 dai (top panel) and 14 dai (bottom panel). Inner plots: Histograms illustrating the distribution of genotypic values for plant development.

**Supplementary Table 1** | Details about the isolation of the strains characterized in this study. CBC: Community-Based Culture.

**Supplementary Table 2** | Primers used to validate the genus level of the OTUs for the informative-driven approach.

**Supplementary Table 3** | *gyrB* primers designed in this study for the informative driven approach.

**Supplementary Table 4** | ANI values among strains within each OTU. Values highlighted in orange indicate that the strain OTU3b_Oxa_1 belongs to a different bacterial species to which belong the strains OTU3a_Oxa_1 and OTU3a_Oxa_2. Values highlighted in light blue indicate the three strains from OTU5 and the six strains from OTU6 belong to two distinct bacterial species. NE: not estimated

**Supplementary Table 5** | Genetic variation among eight *A. thaliana* accessions in response to 22 bacterial strains inoculated at the seed stage in *in vitro* conditions. The statistical term ‘Treatment’ corresponds to the effect of each bacterial strain in comparison with the mock treatment. Bold values indicate significant *p* values after a False Discover Rate (FDR) correction at a nominal level of 5%.

**Supplementary Table 6** | Genetic variation among eight *A. thaliana* accessions in response to 22 bacterial strains inoculated at the seedling stage in *in vitro* conditions. The statistical term ‘Treatment’ corresponds to the effect of each bacterial strain in comparison with the mock treatment. Bold values indicate significant *p* values after a False Discover Rate (FDR) correction at a nominal level of 5%.

**Supplementary Data Set 1.** Taxonomic affiliation after blasting a *gyrB* amplicon of the 12 most prevalent and abundant OTUs against the NCBI database.

**Supplementary Data Set 2.** Forward and reverse *gyrB* primers containing different internal tags in order to multiplex the 7259 amplicons from 76 96-well playes, corresponding to 7259 picked colonies, on a single Miseq run.

**Supplementary Data Set 3.** Barcodes for Illumina libraries with associated SRA numbers.

**Supplementary Data Set 4.** Demultiplexing statistics for each of the 7,259 sequenced colonies.

**Supplementary Data Set 5.** *gyrB* database (v2) used for taxonomic affiliation with DADA2.

**Supplementary Data Set 6.** Taxonomic affiliation of the seven OTUs for which representative strains have been isolated, by using FastANI on the whole-genome sequence of the 20 strains.

