## Supplementary Information for "Investigating genetic diversity within the most abundant and prevalent non-pathogenic leaf-associated bacteria interacting with *Arabidopsis thaliana* in natural habitats"

**Primers design strategy**

All the strains with an identity ≥98% for the *gyrase B* sequence of an OTU of interest were considered as belonging to this OTU. Based on this, we BLASTed each *gyrB* OTU sequence against the gyrB database composed of 38,929 sequences (Supplementary Data Set 5) and retrieved all the *gyrB* sequences with an identity ≥ 85%. Subsequently, we split these sequences in two groups, *i.e.* one group composed of sequences with an identity ≥98% for the OTU of interest (named as OTU sequences) and another group containing all the sequences with an identity between 85% and 98% (named as non-OTU sequences). Then, all the sequences were aligned using MUSCLE program (Edgar 2004) and for each polymorphism along the alignment, an allele frequency was calculated for both group of sequences. We retained all polymorphisms with a difference of allele frequency greater than 90% between the two groups of sequences, thereby maximizing inter-group differentiation while maintaining limiting intra-group diversity. Finally, a combination of 2 positions that allows to be specific to the OTU sequence of interest was selected. Primers were designed in a way that these two selected positions were located at their 3' end, in where mismatches are more likely to have a stronger impact on the specificity. The list of the OTU specific primers are given in Supplementary Table 2.

**Supplementary Figure 1.** CBC and informative-driven approaches to isolate representative strains of the 12 most abundant and prevalent leaf OTUs across 163 natural populations of *Arabidopsis thaliana* located south-west of France.

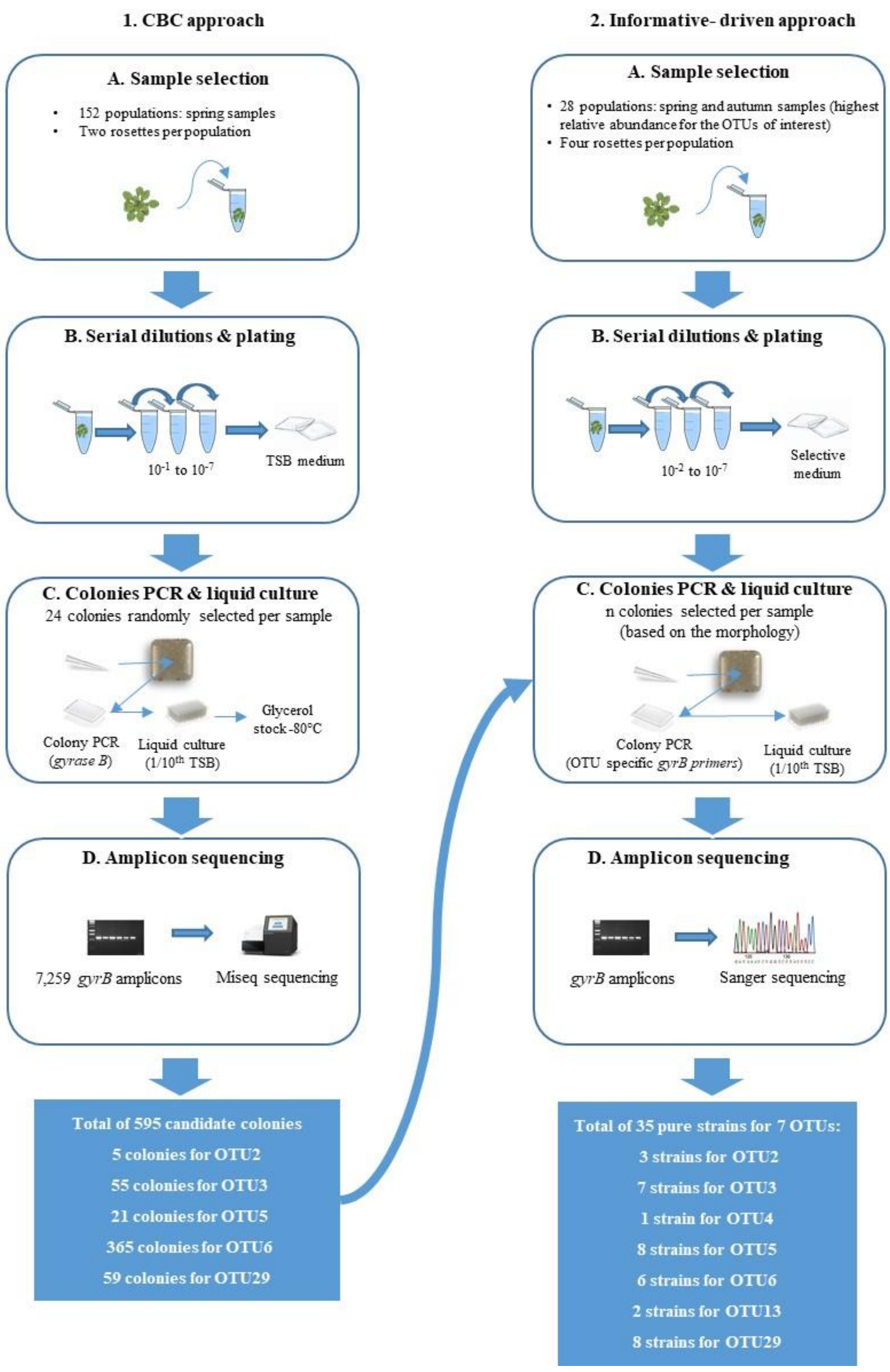

**Supplementary Figure 2.** Pictures illustrating the habitats of the seven accessions chosen to test host genetic variation in response to bacterial isolates.

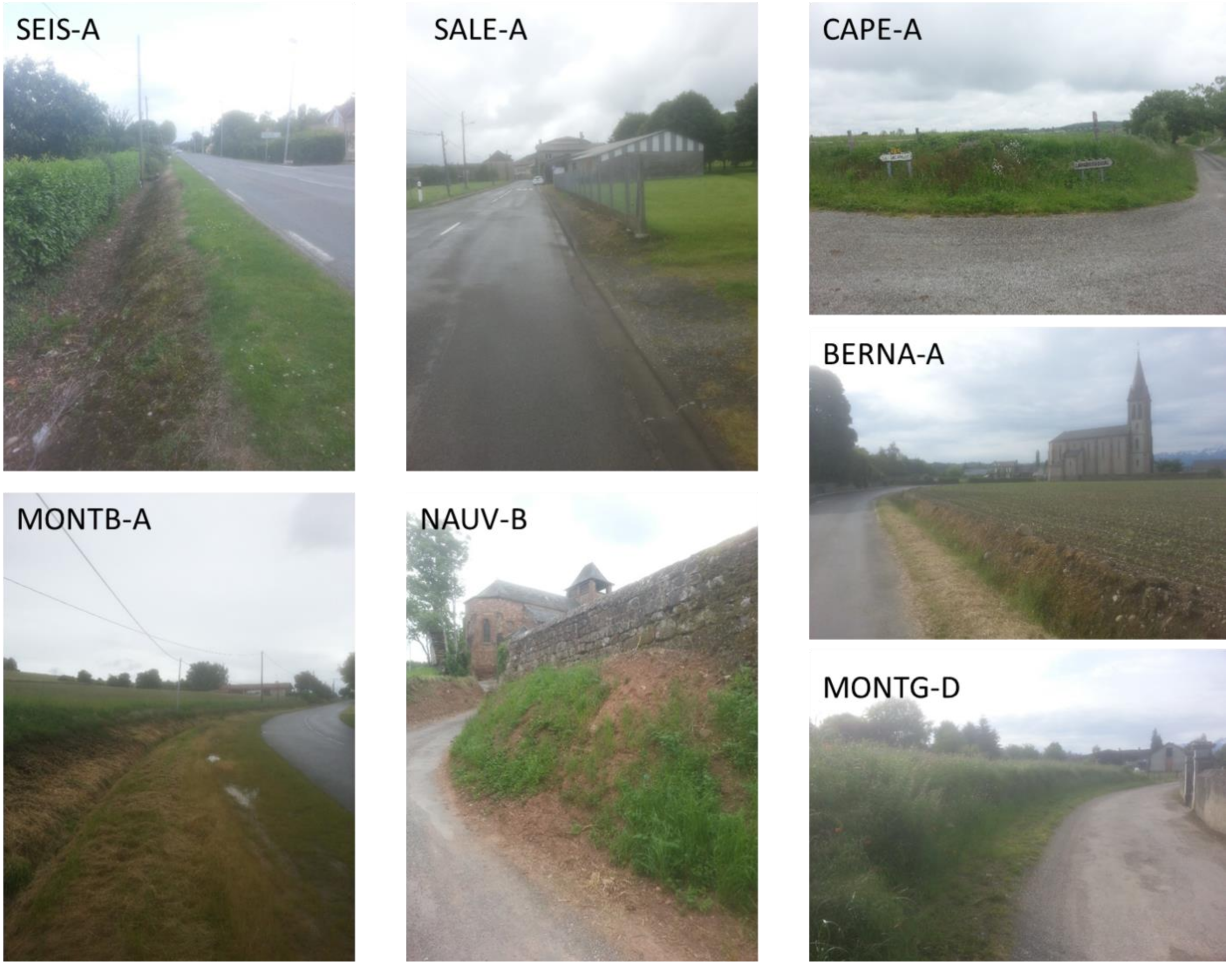

**Supplementary Figure 3.** Intra-OTU genetic variation for growth kinetics on two media with contrasting nutrient availability. **(A)**, **(C)**, **(E)** and **(G)** Intra-OTU genetic variation for the OTUs 4, 5, 13 and 2 on the R2A minimal medium. **(B)**, **(D)**, **(F)** and **(H)** Within-OTU genetic variation for the OTUs 4, 5, 13 and 2 on the TSB rich medium.

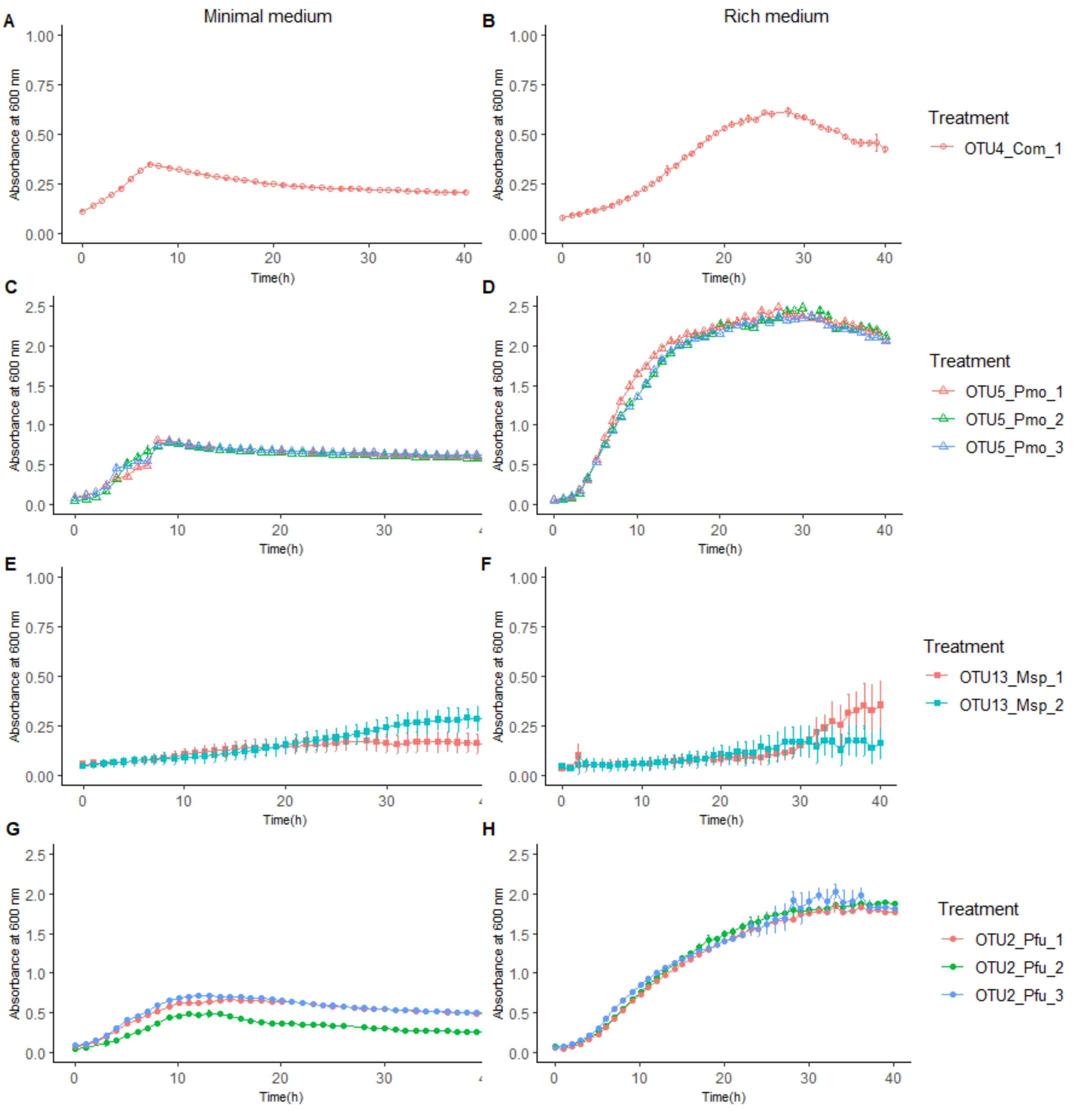

**Supplementary Figure 4.** Double hierarchical clustering based on genotypic values, illustrating the genetic variation among the eight *A. thaliana* accessions in response to the 22 representative bacterial strains belonging to eight OTUs at 28 dai. **(A)** Inoculation at the seed stage with scoring at 14 dai (top panel) and 21 dai (bottom panel). (**B**) Inoculation at the seedling stage with scoring at 7 dai (top panel) and 14 dai (bottom panel). Inner plots: Histograms illustrating the distribution of genotypic values for plant development.

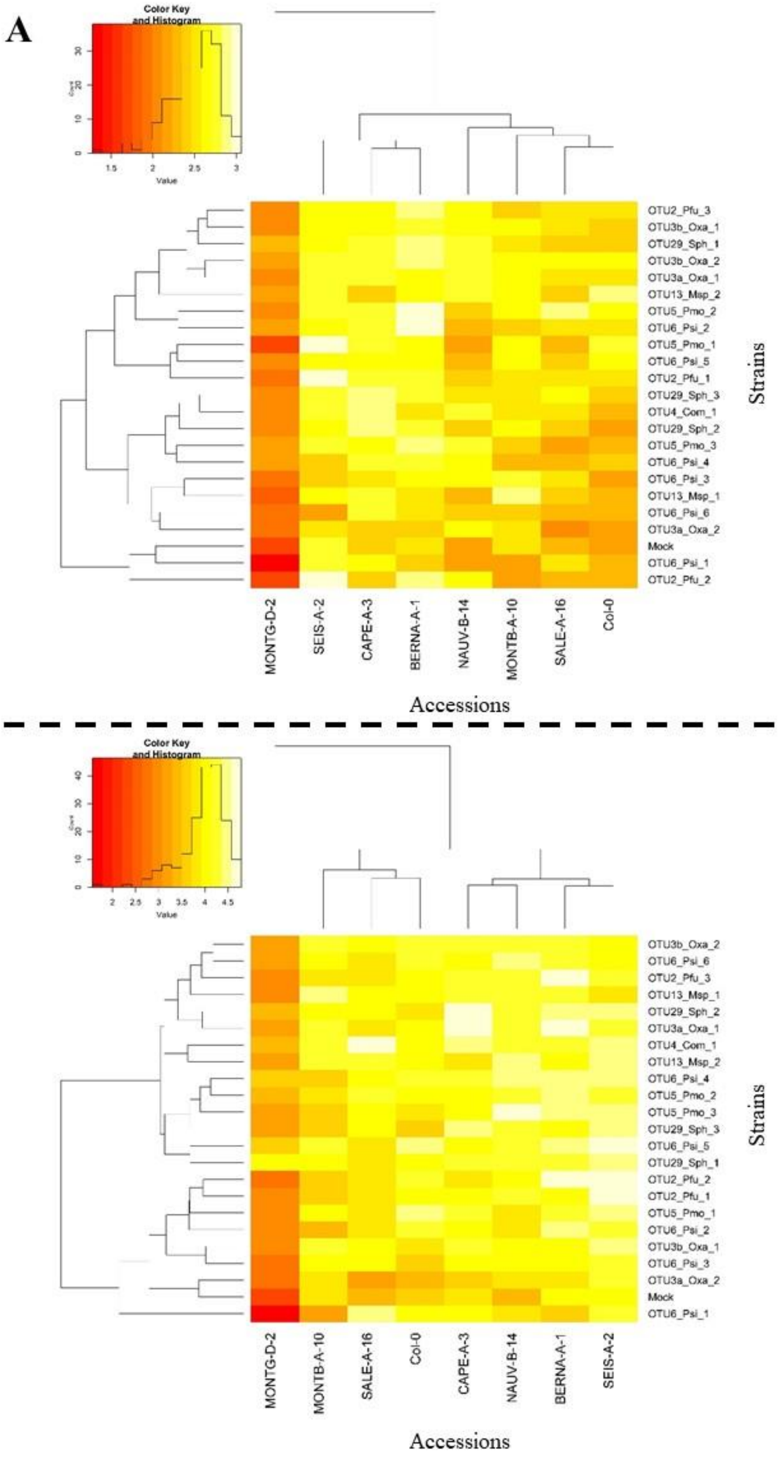

**Supplementary Figure 4 (continued)**

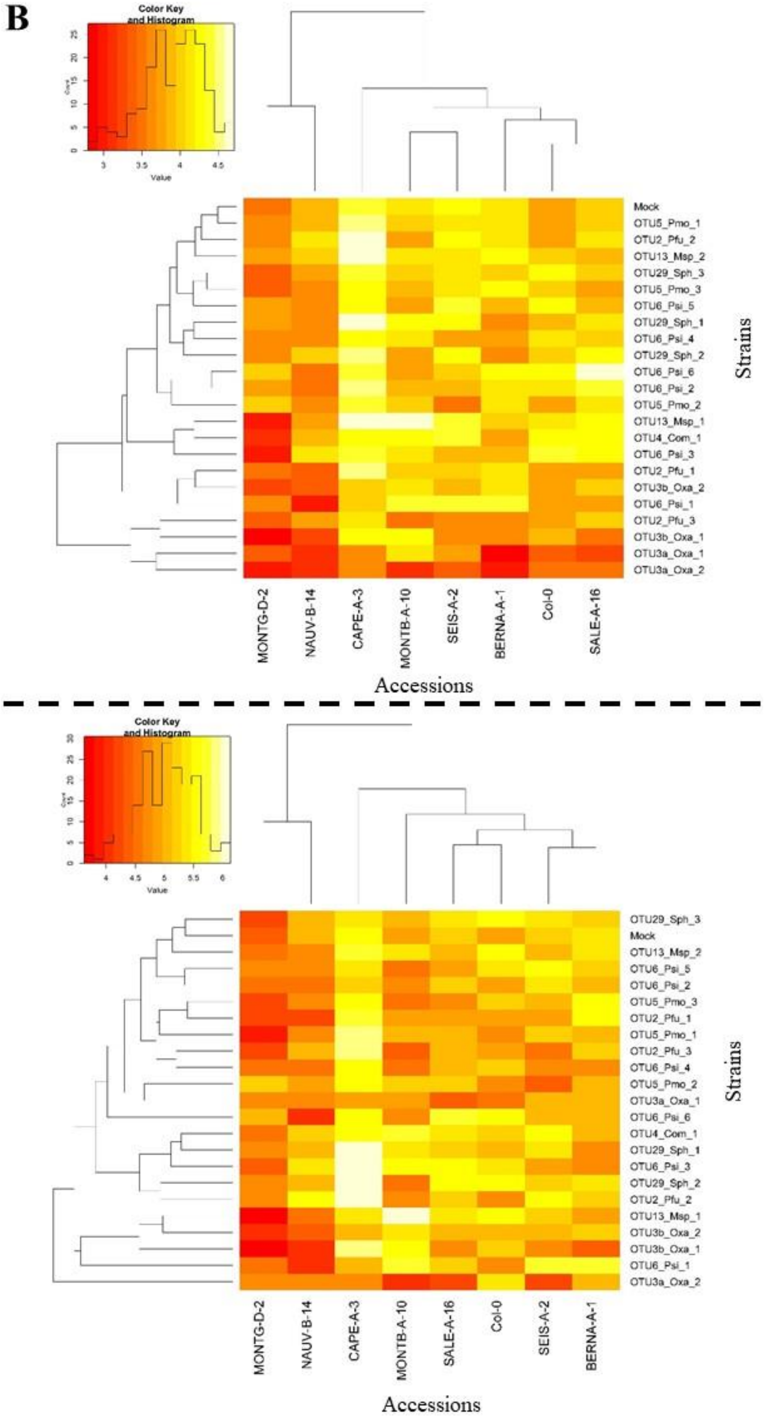

**Supplementary Table 1.** Details about the isolation of the strains characterized in this study. CBC: Community-Based Culture.

| OTU | *gyrB*-based taxonomic affiliation | Approach to isolate strains | Selective media used for OTUs isolation* | Number of pure strains isolated | Growth media for *in planta* experiments | Strains characterized in this study | French location of isolated strains |
|---|---|---|---|---|---|---|---|
| OTU1 | *Sphingomonas* sp. | Informative-driven | R2A (Sigma, France)<br>Nutrient agar (Méridis, France)<br>Streptomycin 100 µg/ml<br>Piperacillin 50 µg/ml<br>(Yim et al., 2010) | 0 | NA | / | NA |
| OTU2 | *Paraburkholderia fungorum* | CBC | TSB (Sigma, France)<br>Polymyxin 50 µg/ml<br>(Loutet et al., 2011) | 3 | TSB | OTU2_*Pfu*_1<br>OTU2_*Pfu*_2<br>OTU2_*Pfu*_3 | Saubens<br>Bagneres-de-Bigorre<br>Bagneres-de-Bigorre |
| OTU3 | *Collimonas* sp. | CBC | R2A | 7 | R2A | OTU3a_*Oxa*_1<br>OTU3a_*Oxa*_2<br>OTU3b_*Oxa*_1<br>OTU3b_*Oxa*_2 | Camarès<br>Réalmont<br>Cintegabelle<br>Saint Angel, Salvagnac |
| OTU4 | *Variovorax* sp. | Informative-driven | R2A<br>Nutrient agar<br>Ampicillin 50 µg/ml<br>(Bergey and Holt, 2000) | 1 | R2A | OTU4_*Com*_1 | Nauviale |
| OTU5 | *Pseudomonas moraviensis* | CBC | TSA (Sigma, France) | 8 | TSA | OTU5_*Pmo*_1<br>OTU5_*Pmo*_2<br>OTU5_*Pmo*_3 | Masseube<br>Villate<br>Barry le Cas (Caylus) |
| OTU6 | *Pseudomonas koreensis* | CBC | TSA | 6 | TSA | OTU6_*Psi*_1<br>OTU6_*Psi*_2<br>OTU6_*Psi*_3<br>OTU6_*Psi*_4<br>OTU6_*Psi*_5<br>OTU6_*Psi*_6 | Espérausses<br>Bagnères-de-Bigorre<br>Mazamet<br>Montiès<br>Banios<br>Monestiès |
| OTU10 | *Acidovorax* sp. | Informative-driven | R2A<br>Nutrient agar | 0 | NA | / | NA |
| OTU12 | *Variovorax* sp. | Informative-driven | R2A<br>Nutrient agar<br>Ampicillin 50 µg/ml | 0 | NA | / | NA |
| OTU13 | *Methylorubrum extorquens* | Informative-driven | R2A<br>Minimum medium supplemented with 120mM of methanol (Guerinot et al., 1990) | 2 | R2A | OTU13_*Msp*_1<br>OTU13_Msp_2 | Castelginest<br>Castelginest |
| OTU14 | *Caulobacter* sp. | Informative-driven | R2A (Sigma, France) | 0 | NA | / | |
| OTU15 | *Modestobacter muralis* | Informative-driven | R2A<br>IPS2 supplemented with nalidixic acid (Qin et al., 2009 ; Golinska et al., 2020), sodium propionate Agar (Qin et al., 2013), Luedemann medium supplemented with nalidixic acid (Qin et al., 2009 ; Luedemann, 1968) | 0 | NA | / | |
| OTU29 | *Sphingomonas melonis* | CBC | R2A (Sigma, France)<br>Nutrient agar (Méridis, France)<br>Streptomycin 100 µg/ml | 8 | R2A | OTU29_*Sph*_1<br>OTU29_Sph_2<br>OTU29_Sph_3 | Martres-Tolosane<br>Saint-Thomas<br>Lamasquère |

* Some strains retrieved from the CBC were not pure and then further purified with selective media.

**Supplementary Table 2.** Primers used to validate the genus level of the OTUs for the informative-driven approach. NA: not available.

| OTU | gyrB-based taxonomic affiliation | Primers ID | Primer sequences | Amplicon size | Annealing temperature | PCR conditions | | | Source |
|---|---|---|---|---|---|---|---|---|---|
| OTU1 | *Sphingomonas* sp. | Sph-spt 295f | CGATCCCTTCGCGATCGTG | 408bp | 55°C | 94°c | 5min | | Yim et al., 2010 |
| | | Sph-spt 713R | TGGCGGAAGCGGACGATC | | | 94°c | 1min | x30 | |
| | | Sph-spt 694f | GAGATCGTCCGCTTCCGC | 289bp | | 55°c | 1min | | |
| | | Sph-spt 983R | CCGACCGATTTGGAGAAG | | | 72°c | 1min | | |
| | | | | | | 72°c | 5min | | |
| OTU2 | *Paraburkholderia fungorum* | gyrBB1f | ACCGGTCTGCAYCACCTCGT | 738bp | 60°C | 94°c | 5min | | Spilker et al., 2009 |
| | | gyrBB2r | YTCGTTGWARCTGTCGTTCCACTGC | | | 94°c | 1min | x30 | |
| | | recAB1f | AGGACGATTCATGGAAGAWAGC | 704bp | 58°C | 60/58°c | 1min | | |
| | | recAB2r | GACGCACYGAYGMRTAGAACTT | | | 72°c | 1min | | |
| | | | | | | 72°c | 5min | | |
| OTU3 | *Collimonas* sp. | NA | | | | | | | / |
| OTU4 | *Variovorax* sp. | VarF | CAATCGTGGGGGATAACGC | 88bp | 60°C | 94°c | 5min | | Bers et al., 2011 |
| | | VarR | GGCCGCTCCATTCGCGCA | | | 94°c | 1min | x30 | |
| | | | | | | 60°c | 1min | | |
| | | | | | | 72°c | 1min | | |
| | | | | | | 72°c | 10min | | |
| OTU5 | *Pseudomonas moraviensis* | gyrB+271ps | TCBGCRGCVGARGTSATCATGAC | 762bp | 62.9°C | 94°c | 5min | | Hwang et al., 2005<br>Gholami et al., 2016 |
| | | gyrB-1022ps | TTGTCYTTGGTCTGSGAGCTGAA | | | 94°c | 1min | x30 | |
| | | rpoD+364s | CAGGTGGAAGACATCATCCCGCAGT | 704bp | 62.2°C | 62.9/55°c | 1min | | |
| | | rpoD-1222ps | GYGAAGGCGARATYGRAATCG | | | 72°c | 30sec | | |
| | | OprI-F | ATGAACAACGTTCTGAAATTCTCTGCT | 249bp | 55°C | 72°c | 5min | | |
| | | OprI-R | CTTGCGGCTGGCTTTTTCCAG | | | | | | |
| OTU6 | *Pseudomonas koreensis* | gyrB+271ps | TCBGCRGCVGARGTSATCATGAC | 762bp | 62.9°C | 94°c | 5min | | Hwang et al., 2005<br>Gholami et al., 2016 |
| | | gyrB-1022ps | TTGTCYTTGGTCTGSGAGCTGAA | | | 94°c | 1min | x30 | |
| | | rpoD+364s | CAGGTGGAAGACATCATCCCGCAGT | 704bp | 62.2°C | 62.9/55°c | 1min | | |
| | | rpoD-1222ps | GYGAAGGCGARATYGRAATCG | | | 72°c | 30sec | | |
| | | OprI-F | ATGAACAACGTTCTGAAATTCTCTGCT | 249bp | 55°C | 72°c | 5min | | |
| | | OprI-R | CTTGCGGCTGGCTTTTTCCAG | | | | | | |
| OTU10 | *Acidovorax* sp. | Oaf1 | GTCGGTGCTAACGACATGG | 550bp | 55°C | 94°c | 5min | | Fontana et al., 2013 |
| | | Oar1 | AGACATCTCCGCTTTCTTTCAA | | | 94°c | 1min | x25 | |
| | | | | | | 55°c | 30sec | | |
| | | | | | | 72°c | 1min | | |
| | | | | | | 72°c | 10min | | |
| OTU12 | *Variovorax* sp. | VarF | CAATCGTGGGGGATAACGC | 88bp | 60°C | 94°c | 5min | | Bers et al., 2011 |
| | | VarR | GGCCGCTCCATTCGCGCA | | | 94°c | 1min | x30 | |
| | | | | | | 60°c | 1min | | |
| | | | | | | 72°c | 1min | | |
| | | | | | | 72°c | 10min | | |
| OTU13 | *Methylorubrum extorquens* | NA | | | | | | | / |
| OTU14 | *Caulobacter* sp. | NA | | | | | | | / |
| OTU15 | *Modestobacter muralis* | NA | | | | | | | / |
| OTU29 | *Sphingomonas melonis* | Sph-spt 295f | CGATCCCTTCGCGATCGTG | 408bp | 55°C | 94°c | 5min | | Yim et al., 2010 |
| | | Sph-spt 713R | TGGCGGAAGCGGACGATC | | | 94°c | 1min | x30 | |
| | | Sph-spt 694f | GAGATCGTCCGCTTCCGC | 289bp | | 55°c | 1min | | |
| | | Sph-spt 983R | CCGACCGATTTGGAGAAG | | | 72°c | 1min | | |
| | | | | | | 72°c | 5min | | |

**Supplementary Table 3.** *gyrB* primers designed in this study for the informative-driven approach. NA: not available.

| OTUs | gyrB-based taxonomic affiliation | Primers ID | Primer sequences (this study, informative driven approach) | Amplicon size | Annealing temperature | PCR conditions |
|---|---|---|---|---|---|---|
| OTU1 | *Sphingomonas* sp. | OTU_1_spe_82_FAM<br>OTU_1_spe_109_Common | GAAGGTGACCAAGTTCATGCTAGCGCTGGCGGGTCAC<br>GACACCGAGCCATCGGGA | 82bp | 66°C | 95°c 1min<br>[95°c 30sec<br>66°c 30sec<br>72°c 30sec] x35<br>72°c 10min |
| OTU2 | *Burkholderia fungorum* | OTU_2_spe_76_F<br>OTU_2_spe_91_R | GAAGGTGACCAAGTTCATGCTATCGACGAAGCGTTGGCG<br>CCGCGTGAATGATCACYTGGATA | 80bp | 65°C a | 94°c 5min<br>[94°c 30sec<br>65°c 30sec<br>72°c 30sec] x30<br>72°c 5min |
| OTU3 | *Collimonas* sp. | NA | | | | |
| OTU4 | *Variovorax* sp. | OTU_4_spe_145_F<br>OTU_4_spe_202_R | GAAGGTGACCAAGTTCATGCTTCACCGACAACGGCCGT<br>CTCGGTCAGCGCGATCTCC | 114bp | 65°C | 94°c 5min<br>[94°c 30sec<br>65°c 1min<br>72°c 30sec] x30<br>72°c 5min |
| OTU5 | *Pseudomonas moraviensis* | NA | | | | |
| OTU6 | *Pseudomonas koreensis* | OTU_6_fwd<br>OTU_6_rv | GAAGGTGACCAAGTTCATGCTGGATGCCACGACCGTTGTCTT<br>GGATGCCACGACCGTTGTCTT | 104bp | 63°C | 94°c 5min<br>[94°c 1min<br>63°c 30sec<br>72°c 30sec] x30<br>72°c 5min |
| OTU10 | *Acidovorax* sp. | OTU10_SpeF25<br>OTU10_SpeR119 | GCACCATTTGGTGTTCGAGG<br>TTGTCGATGACGCTGATGCT | 141pb | 65°C | 94°c 5min<br>[94°c 1min<br>65°c 1min<br>72°c 1min] x30<br>72°c 10min |
| OTU12 | *Variovorax* sp. | OTU12_142 forward<br>gyrBaR353-1 | GCTGACCGAGCTGCA<br>GGAGTTCAGACGTGTGCTCTTCCGATCTACNCCRTGNARDCCDCCNGA | 150bp | 59°C | 94°c 5min<br>[94°c 1min<br>59°c 30sec<br>72°c 30sec] x30<br>72°c 10min |
| OTU13 | *Methylorubrum extorquens* | NA | | | | |
| OTU14 | *Caulobacter* sp. | NA | | | | |
| OTU15 | *Modestobacter muralis* | NA | | | | |
| OTU29 | *Sphingomonas melonis* | NA | | | | |

**Supplementary Table 4.** ANI values among strains within each OTU. Values highlighted in orange indicate that the strain OTU3b_Oxa_1 belongs to a different bacterial species to which belong the strains OTU3a_Oxa_1 and OTU3a_Oxa_2. Values highlighted in light blue indicate the three strains from OTU5 and the six strains from OTU6 belong to two distinct bacterial species. NE: not estimated

| | OTU29_Sph_3 | OTU29_Sph_1 | OTU29_Sph_2 | OTU2_Pfu_3 | OTU2_Pfu_2 | OTU2_Pfu_1 | OTU3a_Oxa_1 | OTU3b_Oxa_1 | OTU3a_Oxa_2 | OTU5_Pmo_3 | OTU5_Pmo_1 | OTU5_Pmo_2 | OTU6_Psi_2 | OTU6_Psi_5 | OTU6_Psi_1 | OTU6_Psi_3 | OTU6_Psi_6 | OTU6_Psi_4 |
|---|---|---|---|---|---|---|---|---|---|---|---|---|---|---|---|---|---|---|
| OTU29_Sph_3 | 100 | | | | | | | | | | | | | | | | | |
| OTU29_Sph_1 | 98.01 | 100 | | | | | | | | | | | | | | | | |
| OTU29_Sph_2 | 98.05 | 98.03 | 100 | | | | | | | | | | | | | | | |
| OTU2_Pfu_3 | NE | NE | NE | 100 | | | | | | | | | | | | | | |
| OTU2_Pfu_2 | NE | NE | NE | 100 | 100 | | | | | | | | | | | | | |
| OTU2_Pfu_1 | NE | NE | NE | 98.53 | 98.53 | 100 | | | | | | | | | | | | |
| OTU3a_Oxa_1 | NE | NE | NE | NE | NE | NE | 100 | | | | | | | | | | | |
| OTU3b_Oxa_1 | NE | NE | NE | NE | NE | NE | 92.96 | 100 | | | | | | | | | | |
| OTU3a_Oxa_2 | NE | NE | NE | NE | NE | NE | 95.69 | 93.11 | 100 | | | | | | | | | |
| OTU5_Pmo_3 | NE | NE | NE | NE | NE | NE | NE | NE | NE | 100 | | | | | | | | |
| OTU5_Pmo_1 | NE | NE | NE | NE | NE | NE | NE | NE | NE | 98.67 | 100 | | | | | | | |
| OTU5_Pmo_2 | NE | NE | NE | NE | NE | NE | NE | NE | NE | 98.45 | 98.39 | 100 | | | | | | |
| OTU6_Psi_2 | NE | NE | NE | NE | NE | NE | NE | NE | NE | 92.87 | 92.84 | 92.79 | 100 | | | | | |
| OTU6_Psi_5 | NE | NE | NE | NE | NE | NE | NE | NE | NE | 92.88 | 92.91 | 92.82 | 97.79 | 100 | | | | |
| OTU6_Psi_1 | NE | NE | NE | NE | NE | NE | NE | NE | NE | 92.92 | 92.93 | 92.85 | 98.37 | 97.80 | 100 | | | |
| OTU6_Psi_3 | NE | NE | NE | NE | NE | NE | NE | NE | NE | 92.85 | 92.90 | 92.82 | 98.43 | 97.81 | 98.36 | 100 | | |
| OTU6_Psi_6 | NE | NE | NE | NE | NE | NE | NE | NE | NE | 92.89 | 92.88 | 92.81 | 97.81 | 97.92 | 97.79 | 97.79 | 100 | |
| OTU6_Psi_4 | NE | NE | NE | NE | NE | NE | NE | NE | NE | 92.86 | 92.86 | 92.82 | 97.82 | 98.03 | 97.83 | 97.78 | 98.02 | 100 |

**Supplementary Table 5.** Genetic variation among eight *A. thaliana* accessions in response to 22 bacterial strains inoculated at the seed stage in *in vitro* conditions. The statistical term 'Treatment' corresponds to the effect of each bacterial strain in comparison with the mock treatment. Bold values indicate significant *p* values after a False Discover Rate (FDR) correction at a nominal level of 5%.

| | **Model terms** | | | | | | | | | | | | | |
|---|---|---|---|---|---|---|---|---|---|---|---|---|---|---|
| | **Accession** | | **Treatment** | | **Time** | | **Accession * Treatment** | | **Treatment*Time** | | **Accession*Time** | | **Treatment*Accession*Time** | |
| **Strains** | *F value* | *P* | *F value* | *P* | *F value* | *P* | *F value* | *P* | *F value* | *P* | *F value* | *P* | *F value* | *P* |
| OTU2_*Pfu* _1 | 18.14 | **0.000115789** | 6.30 | 0.2684 | 44.43 | **0.0001** | 1.38 | 0.22924 | 0.65 | 0.9955 | 0.83 | 0.77775789 | 0.42 | 1 |
| OTU2_*Pfu* _2 | 20.86 | **0.000115789** | 3.22 | 0.715 | 49.08 | **0.0001** | 0.82 | 0.5693 | 1.08 | 0.9955 | 1.15 | 0.61545 | 0.42 | 1 |
| OTU2_*Pfu* _3 | 11.30 | **0.000115789** | 0.57 | 0.8966 | 32.40 | **0.0001** | 3.19 | **0.00785714** | 0.24 | 0.9955 | 1.28 | 0.61545 | 1.62 | 1 |
| OTU3a_*Oxa* _1 | 20.39 | **0.000115789** | 0.11 | 0.8966 | 42.80 | **0.0001** | 1.00 | 0.4479619 | 0.24 | 0.9955 | 1.49 | 0.61545 | 0.11 | 1 |
| OTU3a_*Oxa* _2 | 13.30 | **0.000115789** | 0.02 | 0.8966 | 33.72 | **0.0001** | 2.28 | 0.05096667 | 0.34 | 0.9955 | 0.91 | 0.7552875 | 0.59 | 1 |
| OTU3b_*Oxa* _1 | 12.41 | **0.000115789** | 0.04 | 0.8966 | 36.29 | **0.0001** | 2.31 | 0.05096667 | 0.42 | 0.9955 | 0.86 | 0.77775789 | 0.75 | 1 |
| OTU3b_*Oxa* _2 | 17.87 | **0.000115789** | 0.29 | 0.8966 | 48.09 | **0.0001** | 1.39 | 0.22924 | 0.08 | 0.9955 | 1.15 | 0.61545 | 0.20 | 1 |
| OTU4_*Com* _1 | 16.36 | **0.000115789** | 2.33 | 0.715 | 48.49 | **0.0001** | 2.26 | 0.05096667 | 0.21 | 0.9955 | 1.16 | 0.61545 | 0.44 | 1 |
| OTU5_*Pmo* _1 | 0.60 | 0.48851 | 0.60 | 0.8966 | 29.23 | **0.0001** | 1.64 | 0.1562 | 1.37 | 0.9955 | 1.37 | 0.61545 | 0.89 | 1 |
| OTU5_*Pmo* _2 | 13.55 | **0.000115789** | 0.49 | 0.8966 | 45.41 | **0.0001** | 4.35 | **0.00055** | 0.03 | 0.9955 | 0.56 | 0.9425 | 1.09 | 1 |
| OTU5_*Pmo* _3 | 0.11 | **0.776704762** | 0.11 | 0.8966 | 42.46 | **0.0001** | 3.96 | **0.00132** | 0.10 | 0.9955 | 0.96 | 0.72438667 | 0.88 | 1 |
| OTU6_*Psi* _1 | 37.07 | **0.000115789** | 0.28 | 0.8966 | 37.71 | **0.0001** | 6.23 | **0.00055** | 0.00 | 0.9955 | 2.70 | **0.0154** | 0.83 | 1 |
| OTU6_*Psi* _2 | 13.81 | **0.000115789** | 0.12 | 0.8966 | 13.81 | **0.0001** | 3.02 | 0.010725 | 0.64 | 0.9955 | 0.80 | 0.77775789 | 0.55 | 1 |
| OTU6_*Psi* _3 | 18.67 | **0.000115789** | 0.03 | 0.8966 | 39.90 | **0.0001** | 2.30 | 0.05096667 | 0.09 | 0.9955 | 1.18 | 0.61545 | 0.44 | 1 |
| OTU6_*Psi* _4 | 13.93 | **0.000115789** | 2.30 | 0.715 | 49.20 | **0.0001** | 4.53 | **0.00055** | 0.63 | 0.9955 | 1.22 | 0.61545 | 0.54 | 1 |
| OTU6_*Psi* _5 | 16.66 | **0.000115789** | 0.25 | 0.8966 | 40.44 | **0.0001** | 1.58 | 0.16964444 | 0.73 | 0.9955 | 1.00 | 0.7051 | 0.46 | 1 |
| OTU6_*Psi* _6 | 14.94 | **0.000115789** | 0.04 | 0.8966 | 45.25 | **0.0001** | 2.05 | 0.07852308 | 0.96 | 0.9955 | 1.07 | 0.64849231 | 0.42 | 1 |
| OTU13_*Msp* _1 | 19.56 | **0.000115789** | 0.02 | 0.8966 | 39.83 | **0.0001** | 1.75 | 0.128975 | 0.01 | 0.9955 | 1.59 | 0.61545 | 0.39 | 1 |
| OTU13_*Msp* _2 | 0.39 | 0.9767 | 0.29 | 0.8966 | 46.69 | **0.0001** | 1.82 | 0.11909333 | 0.19 | 0.9955 | 1.12 | 0.61545 | 0.42 | 1 |
| OTU29_*Sph* _1 | 8.68 | **0.000115789** | 0.51 | 0.8966 | 51.68 | **0.0001** | 6.23 | **0.00055** | 0.59 | 0.9955 | 0.50 | 0.9425 | 1.04 | 1 |
| OTU29_*Sph* _2 | 14.87 | **0.000115789** | 0.37 | 0.8966 | 32.68 | **0.0001** | 1.95 | 0.09287143 | 0.64 | 0.9955 | 1.15 | 0.61545 | 1.15 | 1 |
| OTU29_*Sph* _3 | 13.80 | **0.000115789** | 0.20 | 0.8966 | 37.50 | **0.0001** | 3.32 | **0.00623333** | 0.06 | 0.9955 | 0.06 | 0.9425 | 0.74 | 1 |

**Supplementary Table 6.** Genetic variation among eight *A. thaliana* accessions in response to 22 bacterial strains inoculated at the seedling stage in *in vitro* conditions. The statistical term 'Treatment' corresponds to the effect of each bacterial strain in comparison with the mock treatment. Bold values indicate significant *p* values after a False Discover Rate (FDR) correction at a nominal level of 5%.

| | Model terms | | | | | | | | | | | | | |
|---|---|---|---|---|---|---|---|---|---|---|---|---|---|---|
| | **Accession** | | **Treatment** | | **Time** | | **Accession * Treatment** | | **Treatment*Time** | | **Accession*Time** | | **Treatment*Accession*Time** | |
| **Strains** | *F value* | *P* | *F value* | *P* | *F value* | *P* | *F value* | *P* | *F value* | *P* | *F value* | *P* | *F value* | *P* |
| OTU2_*Pfu* _1 | 12.23 | **0.00010476** | 0.94 | 0.457875 | 24.29 | **0.0001** | 1.25 | 0.39937333 | 0.86 | 0.9002125 | 0.56 | 1 | 0.28 | 1 |
| OTU2_*Pfu* _2 | 9.78 | **0.00010476** | 1.63 | 0.35335385 | 28.75 | **0.0001** | 1.25 | 0.39937333 | 1.45 | 0.9002125 | 0.38 | 1 | 0.29 | 1 |
| OTU2_*Pfu* _3 | 7.82 | **0.00010476** | 0.99 | 0.457875 | 18.07 | **0.0001** | 3.16 | **0.00817143** | 0.26 | 0.96173 | 0.60 | 1 | 0.40 | 1 |
| OTU3a_*Oxa* _1 | 4.75 | **0.00010476** | 4.80 | 0.0856 | 24.31 | **0.0001** | 1.21 | 0.4059 | 0.94 | 0.9002125 | 0.47 | 1 | 0.52 | 1 |
| OTU3a_*Oxa* _2 | 3.28 | **0.0019** | 0.28 | 0.69566316 | 22.00 | **0.0001** | 2.49 | 0.03813333 | 1.70 | 0.9002125 | 1.15 | 1 | 0.51 | 1 |
| OTU3b_*Oxa* _1 | 13.43 | **0.00010476** | 1.08 | 0.457875 | 18.48 | **0.0001** | 3.67 | **0.00264** | 0.23 | 0.96173 | 0.66 | 1 | 0.64 | 1 |
| OTU3b_*Oxa* _2 | 9.23 | **0.00010476** | 0.15 | 0.7356381 | 16.28 | **0.0001** | 0.86 | 0.59257 | 0.16 | 0.96548571 | 0.28 | 1 | 0.35 | 1 |
| OTU4_*Com* _1 | 15.32 | **0.00010476** | 6.10 | **0.0418** | 22.66 | **0.0001** | 1.98 | 0.1094 | 0.54 | 0.9002125 | 0.39 | 1 | 0.22 | 1 |
| OTU5_*Pmo* _1 | 10.69 | **0.00010476** | 0.79 | 0.49525882 | 21.91 | **0.0001** | 0.41 | 0.8976 | 0.35 | 0.96173 | 0.45 | 1 | 0.35 | 1 |
| OTU5_*Pmo* _2 | 6.27 | **0.00010476** | 0.01 | 0.9301 | 16.20 | **0.0001** | 4.22 | **0.00055** | 0.07 | 0.9779 | 0.19 | 1 | 0.57 | 1 |
| OTU5_*Pmo* _3 | 11.78 | **0.00010476** | 6.64 | **0.0418** | 25.77 | **0.0001** | 1.03 | 0.47137895 | 1.03 | 0.9002125 | 0.88 | 1 | 0.17 | 1 |
| OTU6_*Psi* _1 | 0.16 | **0.00010476** | 0.16 | 0.7356381 | 0.16 | **0.0001** | 5.65 | **0.00055** | 0.30 | 0.96173 | 0.38 | 1 | 0.43 | 1 |
| OTU6_*Psi* _2 | 9.07 | **0.00010476** | 19.23 | **0.0022** | 31.73 | **0.0001** | 1.07 | 0.47137895 | 1.89 | 0.9002125 | 0.37 | 1 | 0.44 | 1 |
| OTU6_*Psi* _3 | 11.17 | **0.00010476** | 8.25 | **0.0242** | 28.53 | **0.0001** | 2.51 | **0.03813333** | 1.26 | 0.9002125 | 0.34 | 1 | 0.54 | 1 |
| OTU6_*Psi* _4 | 6.58 | **0.00010476** | 13.68 | **0.0044** | 24.45 | **0.0001** | 1.91 | 0.11825 | 0.61 | 0.9002125 | 0.61 | 1 | 0.20 | 1 |
| OTU6_*Psi* _5 | 8.36 | **0.00010476** | 4.01 | 0.08635 | 24.47 | **0.0001** | 2.42 | 0.04004 | 1.02 | 0.9002125 | 0.56 | 1 | 0.15 | 1 |
| OTU6_*Psi* _6 | 9.84 | **0.00010476** | 7.16 | 0.03483333 | 25.13 | **0.0001** | 5.34 | **0.00055** | 0.94 | 0.9002125 | 0.70 | 1 | 0.57 | 1 |
| OTU13_*Msp* _1 | 18.31 | **0.00010476** | 4.14 | 0.0856 | 26.07 | **0.0001** | 4.70 | **0.00055** | 0.66 | 0.9002125 | 0.68 | 1 | 0.47 | 1 |
| OTU13_*Msp* _2 | 9.32 | **0.00010476** | 6.28 | 0.0418 | 25.29 | **0.0001** | 0.61 | 0.78707619 | 0.57 | 0.9002125 | 0.40 | 1 | 0.32 | 1 |
| OTU29_*Sph* _1 | 8.75 | **0.00010476** | 8.95 | 0.02948 | 26.51 | **0.0001** | 1.37 | 0.36181538 | 0.96 | 0.9002125 | 0.41 | 1 | 0.23 | 1 |
| OTU29_*Sph* _2 | 10.00 | **0.00010476** | 0.54 | 0.5676 | 23.83 | **0.0001** | 3.32 | **0.00623333** | 0.65 | 0.9002125 | 1.05 | 1 | 0.41 | 1 |
| OTU29_*Sph* _3 | 12.37 | **0.00010476** | 12.80 | **0.0044** | 27.49 | **0.0001** | 1.04 | 0.47137895 | 1.29 | 0.9002125 | 0.54 | 1 | 0.19 | 1 |
